# Molecular mechanisms and energetics of lipid droplet formation and directional budding^†^

**DOI:** 10.1101/2023.12.04.570036

**Authors:** Fatemeh Kazemi Sabet, Arash Bahrami, Rikhia Ghosh, Bartosz Różycki, Amir H. Bahrami

## Abstract

The formation and budding of lipid droplets (LDs) are known to be governed by the LD size and by membrane tensions in the Endoplasmic Reticulum (ER) bilayer and LD-monolayers. Using coarse-grained simulations of an LD model, we first show that ER-embedded LDs of different sizes can form through a continuous transition from wide LD lenses to spherical LDs at a fixed LD size. The ER tendency to relax its bilayer modulates the transition via a subtle interplay between the ER and LD lipid densities. By calculating the energetic landscape of the LD transition, we demonstrate that this size-independent transition is regulated by the mechanical force balance of ER and LD-tensions, independent from membrane bending and line tension whose energetic contributions are negligible according to our calculations. Our findings explain experimental observation of stable LDs of various shapes. We then propose a novel mechanism for directional LD budding where the required membrane asymmetry is provided by the exchange of lipids between the LD-monolayers. Remarkably, we demonstrate that this budding process is energetically neutral. Consequently, LD budding can proceed by a modest energy input from proteins or other driving agents. We obtain equal lipid densities and membrane tensions in LD-monolayers throughout budding. Our findings indicate that unlike LD formation, LD budding by inter-monolayer lipid exchange is a tension-independent process.

## Introduction

Lipid droplets (LDs) are cellular organelles with a specific structure, which are used by living cells to store and supply lipids for energy and lipid metabolism and membrane synthesis ^1–7^. Previously perceived as passive energy packages, LDs have recently emerged as highly dynamic, mobile organelles that are actively implicated in diseases such as obesity, hepatitis, and cancer ^3,8–10^, and are involved in steroid synthesis and cellular development ^4^. LDs are distinguished from typical organelles by unique features: unlike other organelles with an aqueous interior, LDs enclose an oily compartment; unlike bilayer-bounded organelles with negligible membrane tensions, LDs are surrounded by a lipid monolayer with a relatively high tension ^11^; the initial core of LDs inside the Endoplasmic Reticulum (ER), i.e. the nascent LD, has a typical size of a few tens of nanometers ranging from 30 to 60 *nm* in diameter ^12,13^. These characteristic features have significant implications for the initial formation of symmetric LDs (with respect to the ER-bilayer) and subsequent budding of asymmetric LDs from the ER.

LD formation, including initial nucleation and shape transition of symmetrical LDs, appears to be mainly driven and regulated by membrane tensions. Under excess energy or stress conditions, neutral lipids are synthesized in the inter-leaflet space of the ER membrane ^14^, thereby creating a tense ER-bilayer ^15^. Beyond a concentration threshold, these neutral lipids – initially dispersed between the leaflets of the ER-bilayer – coalesce to form nascent LDs. This demixing process ^1,16^, driven by the reducing contact between the neutral and membrane lipids, helps the ER relax its tense membrane. The LDs, which comprise a hydrophobic core covered by two lipid monolayers ^13–15^, have been experimentally observed in the form of both eye-shaped lenses (simply referred to as LD lenses hereafter) and spherical LDs ^13,16^, suggesting that these structures are stable under certain conditions. Stability analysis of a model LD relates LD morphological transition (from LD lenses to spherical LDs) to the LD size, membrane bending stiffness, and an imbalance between the surface tensions experienced by the ER-bilayer and LD-monolayers ^11^. These ER and LD-tensions, known to be related to membrane lipid density ^18^, thus significantly contribute to forming and maintaining ER-embedded LDs with different shapes. Nevertheless, the molecular details of how tense LDs form from initial tensionless ER-bilayers and the free energy landscape of LD formation with the particular contributions of membrane tension, membrane bending, and line tensions remain unexplained.

LD transport and degradation occur through budding of LDs from the ER toward the cytosol ^19–21^. This directional budding helps LDs reach cytosolic proteins and lipid-releasing enzymes, which are crucial for LD metabolism ^12,17,20^. LDs, emerged from the ER, grow in size leading to larger LDs on the micron scale, which either detach from or remain attached to the ER membrane, the latter being more likely ^17,19,20^. Multiple mechanisms have been hypothesized to drive the directional budding of LDs wherein the required membrane asymmetry between cytosolic and luminal LD-monolayers is imposed by different means: membrane curvature, generated by asymmetric lipid composition ^22–26^ and protein insertion ^12,13^; tension imbalance between the LD-monolayers ^15^; and lipid number asymmetry between the LD-monolayers ^12,27^.

The specific contributions of membrane curvature asymmetry, membrane tension imbalance, and lipid composition asymmetry to LD budding is not clear. Membrane curvature, generated in the cytosolic monolayer by inserting proteins or intrinsically-curved lipids, is shown to assist LD directional budding ^26,28^. Several proteins and enzymes contribute to LD formation and its directional budding by enforcing membrane tension asymmetry, via regulating lipid composition, or like fat-storage-inducing FIT2 and ARF1 through generating asymmetric lipid number ^12,14,29,30^. The particular role of other proteins such as Seipin, an integral membrane protein that forms a scaffold around the cytosolic LD-monolayer, is also not certain. While Seipin was experimentally found to redundantly take part in the LD budding ^31^, more recent experiments and coarse-grained simulations suggest that it effectively induces LD budding ^32^.

Regardless of the driving mechanism, LD budding toward the cytosol requires a continuous area growth in the cytosolic LD-monolayer and a corresponding area reduction in the luminal one. This idea is further supported by the fact that the LD-bounding monolayer likely originates from the ER cytosolic leaflet ^19,33^. The area asymmetry can be generated in different ways such as lipid biosynthesis ^27,34^, fusion of the transport vesicles to the ER ^34^, ER lipid flippase ^35^, conversion of neutral lipids to phospholipids by FIT2^36^, and lipid transfer from the luminal to the cytosolic LD-monolayer by proteins such as FIT2 and ARF1. It is not clear which of these pathways are responsible for creating membrane area asymmetry and what mechanisms are driving them.

LD budding has been explained in terms of a discontinuous second-order transition from symmetric LDs to completely-budded asymmetric LDs, regulated by the LD size, and ER and LD-tensions ^11,28^. Across a wide range of LD sizes, initially stable LDs with large ER-tensions become metastable for smaller tensions and eventually unstable for negligible ratios of ER to LD-tensions, where the fully-budded LD is stable. Like formation of symmetric LDs, the budding and emergence of asymmetric LDs is thought to be governed by membrane tensions.

The small size of nascent LDs, below the standard optical microscopy resolution, and the dynamic nature of their formation, make it difficult to investigate LD formation and budding experimentally. Computer simulation of coarse-grained models with molecular resolution offers an alternative platform to dynamically monitor LD formation and budding and to elucidate mechanistic aspects of LD biogenesis by precise measurement of its mechanical properties ^37,38^. Molecular dynamics simulations have been successfully used to study different aspects of LD formation such as the role of proteins like Seipin ^39^ and Caveolin ^40^, implications of Triglyceride blisters for LD biogenesis ^41,42^, surface properties of LDs ^43^, and LD budding by lipid exchange and LD-tension imbalance ^12,14^. Here, we used coarse-grained simulations to explore mechanistic aspects and energetics of LD shape transition and budding, which are poorly understood despite extensive experimental and modelling studies ^1^.

To understand LD formation, we started with identifying the relevant LD parameters by exploring spontaneous LD formation. Consistent with prior studies, membrane tensions, proportional to the lipid density or area-per-lipid, were found to determine the morphological behavior of LDs ^14,18^. Contrary to earlier understanding ^11^, we observed various LD shapes, including stable spherical LDs and LD lenses, across several LD sizes, even for very small LDs. Based on these findings, we hypothesized that stable symmetrical LDs of various shapes form via a continuous transition driven by ER and LD tensions, irrespective of LD size

To corroborate this hypothesis and to better understand morphological transitions and stability of the symmetrical ER-embedded LDs, we then explored the transition from LD lenses to spherical LDs at fixed LD sizes. By calculating the free energies, we demonstrated that by reducing the ER-tension, a continuous LD transition takes place from early-stage LD lenses with large ER-tensions to spherical LDs with negligible ER-tensions. In line with previous studies ^15,18^, a subtle interplay between the area-per-lipids in the ER and LD membranes was found to control membrane tensions and the continuous LD transition. The same interplay between the areas-per-lipids in the ER and LD membranes was also found to build large membrane tensions in LD monolayers. These findings promote the previous understanding of LD formation based on size-dependent discontinuous LD transitions ^11,28^.

Starting from symmetrical spherical LDs embedded in the ER, we then exchanged lipids from the luminal to the cytosolic LD-monolayer, demonstrating that the area asymmetry created by lipid exchange between the LD-monolayers can act as a novel mechanism for LD budding. Remarkably, we observed that LD budding by lipid exchange is energetically neutral due to the nearly equal areas-per-lipid and resulting equal tensions in LD-monolayers. This intriguing result suggests that LD budding can proceed in a quasi-equilibrium manner through intermediate stable states with a modest energy input. Yet this slight energy input, likely supplied by proteins, is essential to drive the LD budding. In contrast to previous models that described LD budding as a tension-induced discontinuous transition from symmetric to asymmetric LDs ^11,28^, our results unveil a neutral transition from symmetric spherical LDs to fully-budded asymmetric LDs. Thus, we demonstrate that, unlike LD formation, LD budding is not driven by membrane tensions but can rather occur through lipid exchange between the LD-monolayers, driven by a modest energy input.

The shape of the nascent LDs depends not only on the ER and LD-tensions but also on the tension along the contact line of LDs with the surrounding ER-bilayer ^28,44^. We demonstrated that even for the nano-sized LDs neither line tension nor membrane bending significantly contribute to the LD formation. Comparing our simulation results with the theory of membrane elasticity, we found a strong agreement between the LD shapes predicted by the theory and those observed in the simulations.

## Methods

### Coarse-grained simulations

We performed Dissipative Particle Dynamics (DPD) simulations ^45–47^ of a coarse-grained model to study LD shape transition and budding. Our simulation method has been widely used to study various aspects of both artificial membranes ^48^ and cellular membranes, including membrane fusion ^49^, cellular membrane rupture and damage ^45^, membrane domain simulation ^50^, vesicle dynamics in shear flow ^51^, and membrane-cholesterol interactions ^52^. The soft molecular interactions of DPD allow faster dynamics with larger time steps ^46,53,54^. Therefore, DPD has been particularly effective in reproducing mechanical properties of lipid membranes ^55,56^ such as membrane tensions ^57–59^ as well as the phase behavior of membranes ^46,52^, which mainly contribute to LD formation and budding.

Our coarse-grained LD model is composed of lipid membranes representing the ER-bilayer and LD-monolayers, an aqueous phase representing the luminal and cytosolic liquids, and an oil phase that represents neutral lipids i.e. the fatty core of the LDs. We note that both cytosolic and luminal liquids are taken to be identical, simply represented by water beads in our model. Coarse-grained phospholipids consist of three hydrophilic head-group beads H and two hydrophobic chains each having six tailgroup beads T, see supplementary Fig. S1a. Both cytosolic and luminal aqueous liquids are composed of water beads W. Each neutral lipid is modelled as a single chain consisting of four oil beads O, see supplementary Fig. S1b. All simulations were performed in LAMMPS simulation package ^60^.

All different coarse-grained bead types H, T, W, O have a diameter of *d* ≈ 0.8 *nm*, which defines the length scale in our simulations ^61^. These beads interact by a conservative DPD force

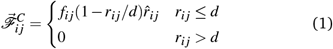

where *r*_*i j*_ is the distance between two beads, *i* and *j*, and the unit vector 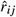 connects the bead *i* to the bead *j*. Whereas the interaction force parameter *f*_*i j*_ varies for different pairs of beads, random and dissipative DPD pairwise forces are the same for all bead pairs ^62^.

In our model, the interaction force parameters between identical pairs of hydrophobic beads O and T were taken to be the same *f*_*OO*_ = *f*_*TT*_ . To induce better demixing of oil molecules from lipid tails and avoid dispersed oils in the ER bilayer, we used a slightly larger interaction force parameter *f*_*OT*_ *> f*_*OO*_ = *f*_*TT*_ between oil and tail beads than those between identical beads. Throughout the manuscript, we used fixed pairwise interaction force parameters *f*_*i j*_ between bead types *i, j* as listed in supplementary Table S1. These molecular interactions are known to represent membrane properties ^61^. Our energy scale is the thermal energy *k*_*B*_*T*, where *k*_*B*_ is the Boltzmann constant and *T* = 298 *K* is the room temperature.

The basic time scale *τ* of the model was taken to be 1 *ns* based on the experimentally measured properties of lipid bilayers, see ref. 60 and the references therein. All DPD simulations were performed using a constant time step of Δ*t* = 0.01*τ* = 0.01 *ns*.

In addition to non-bonded interactions between the beads, we introduced intermolecular bonding and bending forces within the lipid and oil chains. A harmonic bond potential

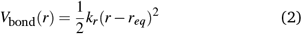

was applied between the adjacent beads of oil and lipid chains at a distance *r* with an equilibrium bond length *r*_*eq*_= 0.5*d* and a bond coefficient *k*_*r*_ = 128 *k*_*B*_*T/d*^2^. Moreover, a bending potential

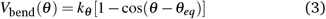

with bending constant *k*_*θ*_ = 15 *k*_*B*_*T* and equilibrium tilt angle *θ*_*eq*_ = 0, acts on all there consecutive beads in each hydrophobic lipid tail and oil chain.

For all simulations performed here, DPD beads were placed in periodic simulation boxes whose dimensions were chosen in a way to give a bead density 3*/d*^3^ = 5.86*/nm*^3^. This density is shown to closely estimate water compressibility at room temperature when the interaction parameter *f*_*WW*_ = 25 *k*_*B*_*T/d* is applied between water beads. We performed three main classes of simulations: spontaneous formation of LDs from initial oil slabs with various values of area-per-lipid, morphological transition of LD lenses to spherical LDs, and LD budding toward the cytosol by inter-monolayer lipid exchange.

## Results and discussion

LD structural biogenesis usually occurs in four successive steps: LD nucleation or coalescence i.e. forming nascent symmetrical LD lenses from initially dispersed neutral lipids in the ER bilayer; LD shape transition from nascent LD lenses to symmetrical spherical LDs; LD budding into the cytosol; and LD fission or detachment from the ER. Our focus, here, is on the two intermediate steps of LD shape transition and budding, before which we identify relevant LD parameters by studying spontaneous LD formation from initial slabs.

LDs, considered here, are either embedded in or attached to the ER membrane and consist of two distinct luminal and cytosolic LD-monolayers, each continuous with its corresponding ER leaflet. Although our membranes are much simpler than actual ER and LD membranes, the lateral bilayer and LD-surrounding monolayers as referred to as the ER-bilayer and LD-monolayers, hereafter. We employed coarse-grained simulations of an LD model to identify LD parameters via spontaneous formation of different LD shapes, and to study LD transition from LD lenses to spherical LDs, and LD budding to the cytosol.

### Relevant LD parameters

LD formation, triggered by coalescence of initially dispersed neutral lipids (referred to as oil hereafter) between the ER leaflets, can be understood using the theoretical framework of ternary phase diagram for surfactant-oil-water mixtures ^15,29^. Within this framework, LD formation is driven by the reducing contact area between oil and aqueous phases, which lowers the energetic contribution of membrane tensions.

ER-attached LDs consist of two membrane interfaces: the ER-bilayer that separates the two aqueous media—the cytoplasm and the ER lumen; and the two LD-monolayers each separating oil from an aqueous medium. Due to oil hydrophobicity, LD-tensions (at the oil-water interface) are generally larger than the ER-tension (at the water-water interface). As a result, LDs naturally tend to reduce the surface area of their LD-monolayers and expand that of the ER-bilayer. Therefore, the initial state of fully-spread neutral lipids between the ER-leaflets with maximum monolayer area—best represented by a slab structure— spontaneously transforms to an LD lens with smaller monolayers and a larger bilayer. The slab-to-lens transition provides a proper platform to identify relevant LD parameters.

The slab consists of two flat monolayers which are exposed to water from above and below and cover a central oil slab with their hydrophobic chains facing the oil, see Fig. 1a. Slab monolayers are made from *N*_*lip*_ lipids, equally partitioned between the two monolayers of a fixed width *L*. The slab-to-lens transition and the resulting LD shapes depend on four parameters: The size of the droplet i.e. the oil volume; the size of the membrane i.e. the area *A*_*s*_ of the slab; and the bilayer and monolayer tensions.

**Fig. 1.**
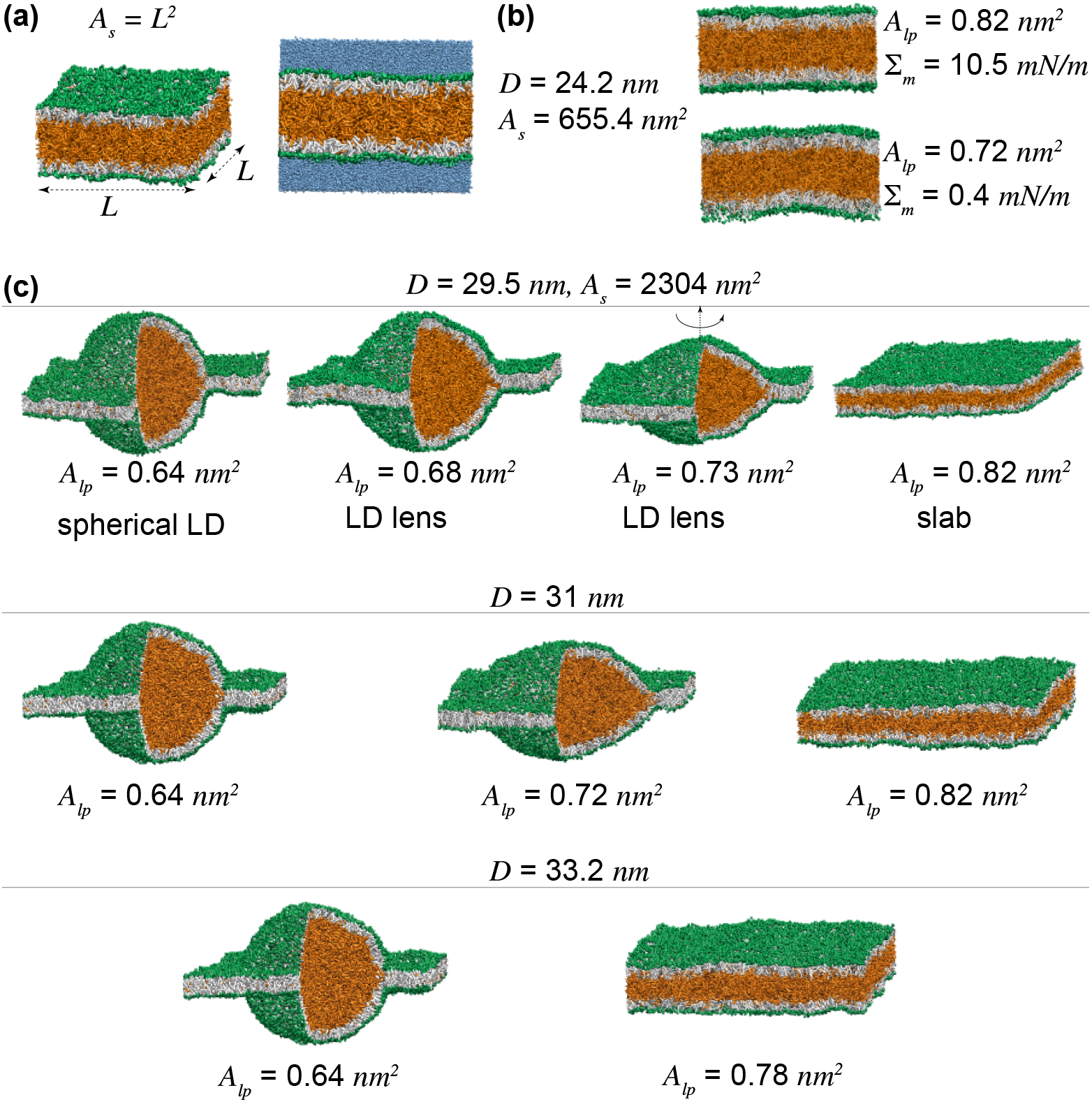
LD parameters. (a) Initial slab morphology of width *L* with surrounding water beads (blue). (b) Two slabs of size *D* = 24.2 *nm* with different projected areas-per-lipid, *A*_*lp*_, and corresponding monolayer tensions. (c) LD shapes including slabs, spherical LDs, and LD lenses, spontaneously formed from initial slabs of various sizes with different values of LD parameters, *D* and *A*_*lp*_.

The size of an LD is given by the volume *V* of its oil content, that is the volume of the neutral lipids. In our model, the size of every LD structure, including different LD lenses and the initial slab configuration, with a given oil volume *V* is defined as the external diameter 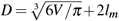 of a spherical monolayer enclosing the same oil volume, where *l*_*m*_ is the thickness of the LD-monolayer.

The area *A*_*s*_ of the slab is defined as the surface area of one of the equal flat monolayers of the initial slab. The slab area *A*_*s*_ = *L*^2^, together with the LD size *D*, determine the surface areas of the ER-bilayer and LD-monolayers of the resulting LDs. For a given *D*, the minimum slab area, *A*_*sm*_ = *πD*^2^*/*4 + *L*^2^, is required to form spherical LDs in a box of width *L*, below which initial slabs retain their flat shape, irrespective of the ER and LD-tensions.

For a given lipid composition, membrane tension has been shown to be directly proportional to its area-per-lipid, for both lipid bilayers and monolayers ^61,63^. The area-per-lipid is defined as the average surface area occupied by one lipid molecule. Each slab with a given number of lipids *N*_*lip*_ has a fixed projected area-per-lipid *A*_*lp*_ = 2*A*_*s*_*/N*_*lip*_, defined as the average area-per-lipid projected into the horizontal plane. The slab monolayer tension ∑_*m*_ is proportional to *A*_*lp*_, see Fig. 1b for small slabs with *D* = 24.2 *nm* and *A*_*s*_ = 655.4 *nm*^2^. Similar behavior was observed for flat bilayers, see supplementary Fig. S2 for more details. The bilayer and monolayer tensions can be thus replaced by their areas-per-lipid. For a given area *A*_*s*_, each slab is characterized by two independent parameters: The LD size *D*, and the projected area-per-lipid *A*_*lp*_. These parameters determine the resulting LD structure that spontaneously forms from the initial slab.

To see how these parameters affect LD shapes, we simulated slabs of a fixed area *A*_*s*_ = 2304 *nm*^2^ and different LD sizes *D* = 29.5, 31, and 33.2 *nm*. For a given LD size *D*, the initial flat monolayers with a small *N*_*lip*_ corresponding to a large *A*_*lp*_, maintained their flat shapes stabilized by large monolayer tensions, see the right snapshots in Fig. 1c. Monolayers with more lipids i.e. a smaller *A*_*lp*_, spontaneously transformed into LD lenses as seen in the middle columns in Fig. 1c. Further increase in the number of lipids of the initial slab eventually led to the spontaneous formation of spherical LDs with smallest *A*_*lp*_’s. LD lenses and spherical LDs that enclose a central axisymmetric oil compartment have been also observed experimentally ^1,16^.

Similar to previous studies ^11,14^, our results indicate that the LD shape is related to its size and ER and LD values of areaper-lipid, and thus ER and LD-tensions. Theoretical models have linked the shape of the symmetrical LD lenses to the LD size ^11^. For a given tension imbalance between the ER and LD membranes, spherical LDs only formed for a relatively large LD volume making LD formation a size-dependent process. However, we observed nano-sized spherical LDs for sufficiently small values of *A*_*lp*_. Even smaller LDs were detected in multiple spherical structures we obtained for small values of *A*_*lp*_. Some of these LDs are shown in Fig. S3. We also observed all LD shapes for three different nano-sized droplets we simulated here. Therefore, unlike the conventional view, our results suggest size-independent transition pathways of different character between the LD shapes.

To better understand LD formation, we explored the hypothesis that symmetrical spherical LDs can form via a size-independent continuous transition from LD lenses driven by energetic forces of ER and LD-tensions. Starting from an LD lens of size *D* = 29.5 *nm*, spontaneously formed from a slab, we changed the box width *L* to study LD shape transition in a systematic manner. By calculating membrane tensions and the corresponding *A*_*l*_ ‘s in the ER-bilayer and LD-monolayers, we explored the transition from nascent LD lenses to spherical LDs and revealed how the subtle interplay between the ER and LD areas-per-lipid govern this transition and lead to tense LD-monolayers and nearly tensionless ER-bilayer. See Methods and Supplementary Information for more details on the simulation protocols.

### Tension-induced transition of LD lenses to spherical LDs

Each LD lens is composed of two symmetrical monolayers connected to the lateral ER-bilayer. The shape of an LD lens of size *D* is characterized by the lens angle *θ* obtained from fitting two spherical caps to its cytosolic and luminal monolayers, see Fig. 2a.

**Fig. 2.**
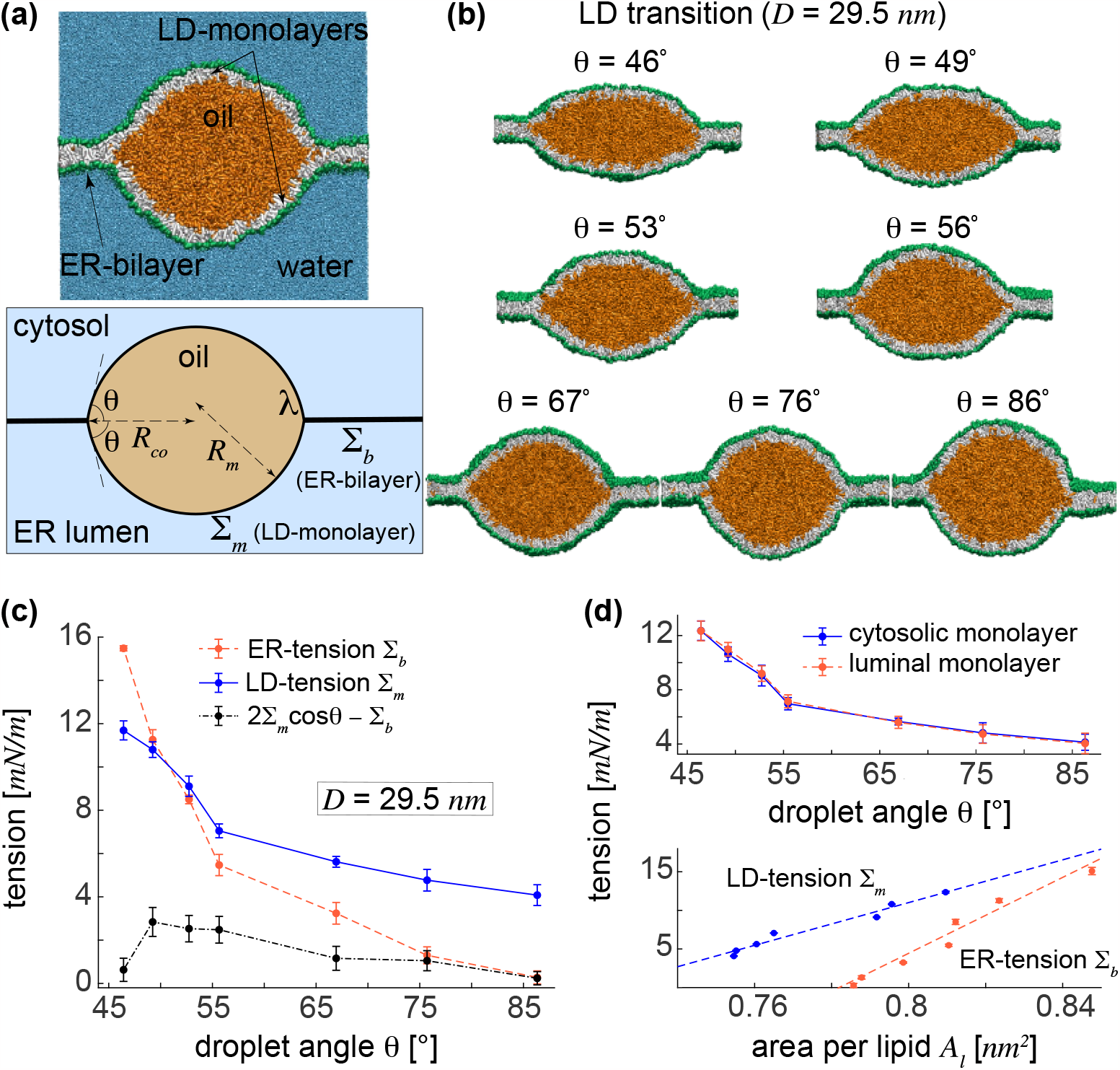
Tension-induced transition of LD lenses to spherical LDs. (a) A model LD and a schematic symmetrical LD with LD angle *θ* . (b) Morphological transition of an initial LD lens of size *D* = 29.5 *nm* to an almost spherical LD. (c) ER and LD-tensions for the LD of size *D* = 29.5 *nm*. (d) Equal tensions in the cytosolic and luminal LD-monolayers verify our accurate tension calculation. Membrane tensions are proportional to *A*_*l*_ for both the ER-bilayer and LD-monolayers.

We obtained different LD lenses of a fixed size *D* = 29.5 *nm* with lens angles in the range 46^*°*^ *< θ <* 86^*°*^, each stabilized for a given box width *L* with the corresponding *A*_*lp*_, implying the stability of intermediate lenses, see Fig. 2b.

For each intermediate lens with angle *θ*, we calculated individual ER and LD-tensions, ∑_*b*_ and ∑_*m*_, respectively. The results are shown in Fig. 2c. The LD-tension was taken to be the average ∑_*m*_ = 0.5(∑_*cm*_ + ∑_*lm*_) of its almost equal components in the cytosolic and luminal monolayers of the symmetrical LD, see the top panel in Fig. 2d. For the wide LD lens with the smallest angle *θ* = 46^*°*^, we found large ER and LD-tensions of ∑_*b*_ = 15.5 *mN/m* and ∑_*m*_ = 11.7 *mN/m*, with the ER-bilayer being more tense than the LD-monolayers. LD lenses in boxes of smaller width *L* and thus smaller *A*_*lp*_’s exhibited lower ER and LD-tensions. Upon approaching more spherical lenses with larger lens angles, the initially tense ER-bilayer exhibited greater relaxation of tension compared to the curved LD-monolayers resulting in a tension crossover, at about *θ* = 50^*°*^ for an LD size *D* = 29.5 *nm*, beyond which the ER-tension fell below the LD-tension. The same trend persisted for larger lens angles leading to an almost tensionless ER-bilayer (∑_*b*_ ≈ 0) for nearly spherical LDs with *θ >* 85^*°*^, wherein LD-monolayers maintained a relatively large tension of about 4 *mN/m*. To understand this behaviour and reveal its underlying molecular mechanism, we focused on the molecular structures of ER and LD membranes by calculating their actual areas-per-lipid *A*_*l*_ .

The spontaneous formation of LDs from initial slabs was observed to depend on the projected area-per-lipid *A*_*lp*_, which is related to the ER and LD-tensions of the final LDs. Membrane tensions are, however, proportional to *A*_*l*_ values in the ER-bilayer and the LD-monolayers. Upon decreasing *L*, the decrease in *A*_*lp*_ is nearly equally shared between the flat ER-bilayer and the curved LD-monolayers as reflected in *A*_*l*_ ‘s of ER and LD membranes, see the bottom panel of Fig. 2d. Both ER and LD-tensions linearly decrease with reducing areas-per-lipid. The slope of the fitted line is proportional to the bending rigidity of the membrane and is thus nearly two folds larger for the ER-bilayer compared to the LD-monolayers. It is indeed this difference in the slope of the linear relations between *A*_*l*_ ‘s and membrane tensions in the ER-bilayer and LD-monolayers that gives rise to the stronger drop of the ER-tension compared to LD-tension and results in tensionless ER-bilayer and tense LD-monolayers of spherical LDs. The bottom panel of Fig. 2d displays the fitted lines to the ER and LD-tensions versus their corresponding *A*_*l*_ for an LD lens of size *D* = 29.5 *nm*. As the shape of the LD lens approaches a sphere, *A*_*l*_ decreases about 0.06 *nm*^2^ in both the ER-bilayer and LD-monolayers. As a result, membrane tension drops more strongly in the ER-bilayer than in the LD-monolayers eventually leading to a nearly tensionless ER with ∑_*b*_ = 0.3 *mN/m* and a tense LD with ∑_*m*_ = 4.1 *mN/m* for the almost spherical LD with *θ* = 86^*°*^.

In the absence of the ER-bilayer and LD-monolayers at the liquid interfaces, the ternary phase model relates interfacial tensions at oil-water and water-water interfaces from the force balance equation 2∑_*OW*_ cos *θ* − ∑_*WW*_ = 0. In their presence, however, this relation is slightly shifted resulting in a tension difference Δ∑ = 2∑_*m*_ cos *θ* − ∑_*b*_ *>* 0 for LDs as shown by the black dash-dotted line in Fig. 2c. This slight deviation is presumably due to the negligible contributions from LD-monolayers bending and the line tension. The tension difference Δ∑ is proportional to the driving force which pulls the ER bilayer and transforms LD lenses to the spherical LDs. Negligible values of Δ∑ indicate that LD transition is primarily regulated by the force balance or mechanical equilibrium which is independent from the LD size. Assuming Δ∑ = 0, the crossover point of the ER and LD-tensions ∑_*b*_ = ∑_*m*_ is obtained as *θ* = 60^*°*^ for all LD sizes. For initial wide lenses with sufficiently small *θ*, the ER-tension is in the order of 2∑_*m*_ and thus larger than the LD-tension. During the transition from LD lenses to spherical LDs, the luminal and cytosolic LD-tensions remained almost identical, as expected due to the vertical symmetry of the LD lens, indicating the accuracy of our tension calculations, see the top panel in Fig. 2d.

We have demonstrated quantitatively that the subtle interplay between *A*_*l*_ values in the ER and LD membranes underlies the continuous LD transition from wide LD lenses to spherical LDs via intermediate LD lenses of different widths. This interplay between the ER and LD areas-per-lipid adjusts ER and LD-tensions and fulfils the force balance independent from the LD size. The slight deviation from this balance provides the driving force for the LD transition. While initial wide LD lenses with sufficiently small angles have larger ER than LD-tensions, a subtle sharing of *A*_*l*_ leads to larger LD than ER tension beyond a crossover point, resulting in large LD and negligible ER-tensions for nearly spherical LDs. This trend, observed here for nano-sized LDs, thus appears to be valid for different LD sizes resulting in a continuous, size-independent LD transition. The LD transition thus resembles a second-order phase transition implying the stability of all LD lenses with different values of *θ*, consistent with experimental observations ^12,14^. Our results are consistent with recent theoretical models that explain the transition from LD lenses to spherical LDs in terms of the relative ratio of ER and LD-tensions, where substantially larger ER-tensions—compared to LD-tensions—stabilized LD lenses ^11,15^. Contrary to the conventional view, we find that the LD transitions is continuous and does not depend on the LD size.

### LD shapes from membrane elastic model

Shape transformations of LDs can also be understood on the basis of membrane elasticity theory. Unlike the simplified spherical cap model that assumes LD-monolayers as two spherical caps, see the bottom panel in Fig. 2a, the elastic model considers them as smooth surfaces whose shapes are found from minimizing the free energy of the ER-LD system. See the Supplementary Information for more details about the elastic model. Here, we combined this model with the spherical cap model to verify our simulation results, in particular the line tensions we computed for LDs. We focused on the same LD of size *D* = 29.5 *nm* discussed before. For each LD lens in Fig. 2b, we used ER and LD-tensions, ∑_*b*_ and ∑_*m*_, obtained from molecular simulations, to calculate the line tension *λ* from the spherical cap model. We then used these tensions to find the corresponding LD shapes from the elastic membrane model. Remarkably, we found a strong agreement between the LD profile (yellow curves) and the outer surface of neutral lipids from the coarse-grained simulations for different LD angles, see Fig. 3. Using the theoretical framework of the elastic membrane model, we thus validated the membrane and line tensions, ∑_*b*_, ∑_*m*_, and *λ*, obtained from molecular simulations and the spherical cap model. The line tensions we calculated here are also close to those recently found for aqueous droplets in contact with membranes ^57,59^.

**Fig. 3.**
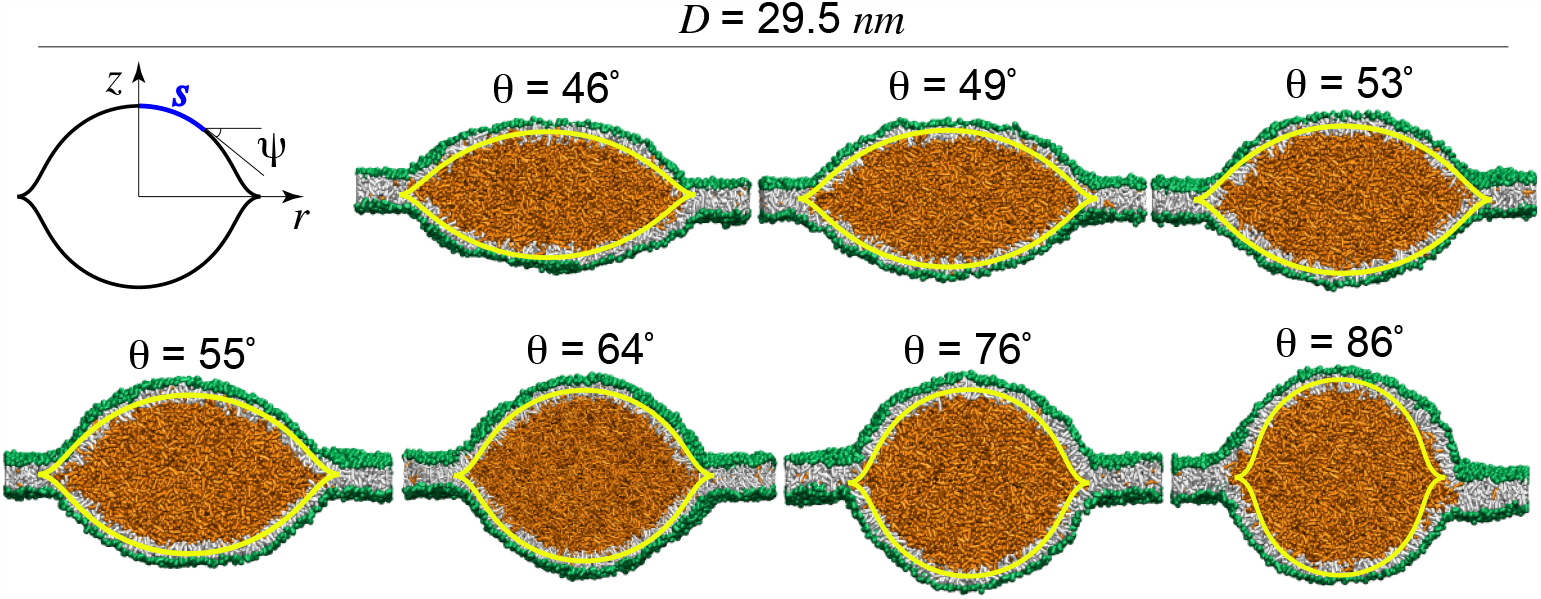
Comparing LD shapes from coarse-grained simulations and membrane elastic model. Symmetrical LD-lenses of size *D* = 29.5 *nm* with different lens angles from molecular simulations, shown in Fig. 2b, with membrane profiles from elastic membrane model (yellow curves). Membrane profiles are parametrized with the tangent angle *ψ* = *ψ*(*s*) as a function of the arc length *s* as shown in the top left panel.

### LD budding to the cytosol

We have understood, thus far, how initially-nucleated LD lenses continuously transform to spherical LDs. Next, we will explore the possibility of LD budding and emergence to the cytosol by exchanging lipids from the luminal to the cytosolic LD-monolayer and examine how membrane tensions and areas-per-lipid vary during the LD budding. The asymmetry in the lipid number is quantified by the rescaled difference Δ = (*N*_*cm*_ − *N*_*lm*_)*/N*_*lip*_ between the number of lipids in the cytosolic and luminal LD-monolayers, *N*_*cm*_ and *N*_*lm*_, respectively. This asymmetry is created by taking lipids from the luminal monolayers into the cytosolic one at constant total number of lipids *N*_*lip*_. We note that the lipid exchange in nature proceeds via a non-equilibrium process. To calculate the free energies of LD budding, however, we equilibrated each LD with a given Δ.

Starting from the spherical LD of size *D* = 29.5 *nm*, we gradually relocated lipid molecules from the luminal monolayer to the cytosolic one, resulting in continuous increase of Δ and subsequent LD budding into the cytosol. Upon increasing Δ, we observed an asymmetric LD structure characterized by two spherical caps as the cytosolic and luminal monolayers with monolayer angles, *θ*_*c*_ and *θ*_*l*_, respectively. As Δ was increased, *θ*_*l*_ kept decreasing while the cytosolic monolayer bulged out by increasing *θ*_*c*_ until the luminal monolayer became relatively small (Fig. 4a), corresponding to the almost fully-budded LD with *θ*_*c*_ = 163^*°*^.

**Fig. 4.**
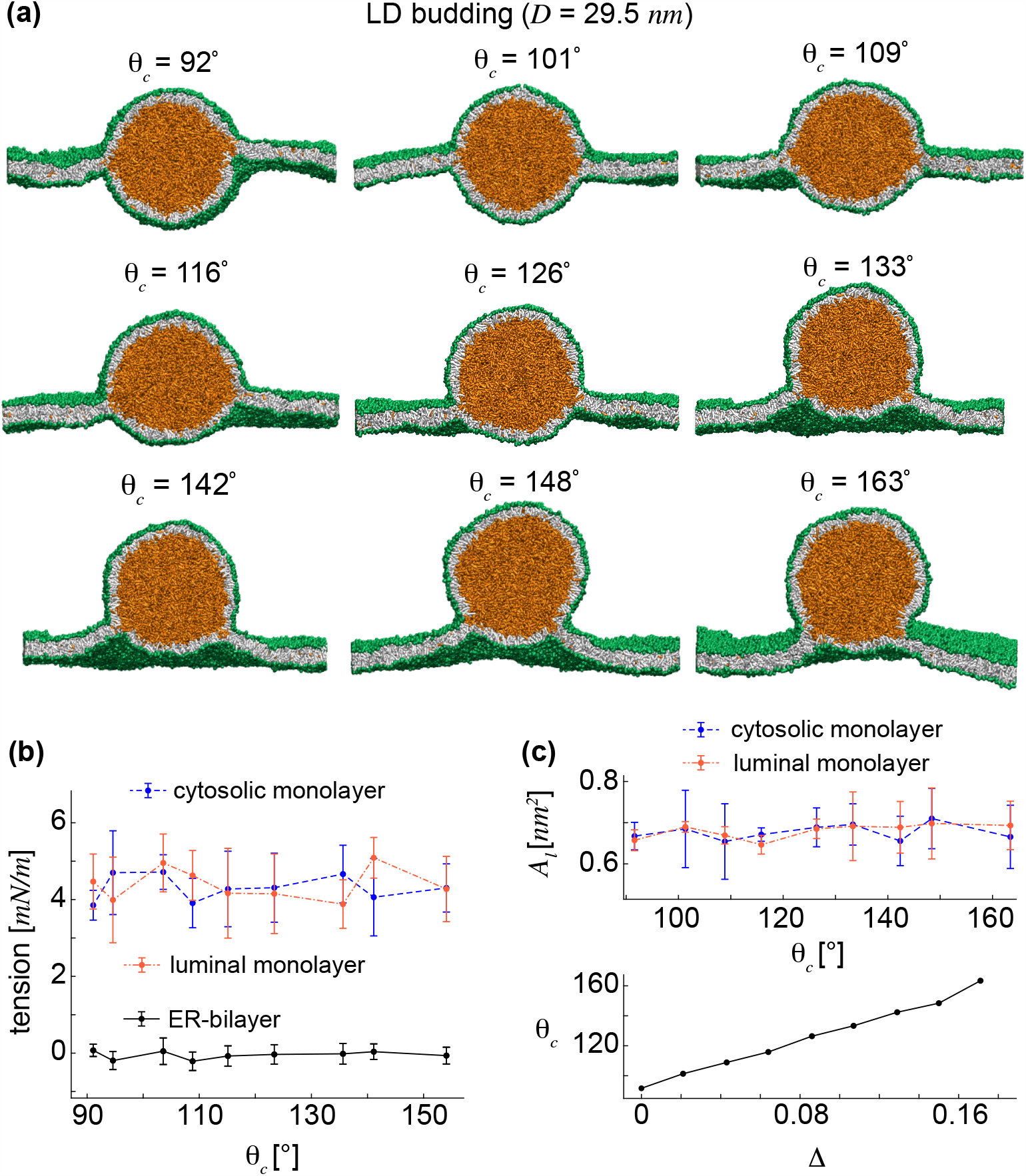
Directional LD budding into the cytosol by inter-monolayer lipid exchange. (a) Schematic LDs budding to the cytosol by lipid exchange with the corresponding cytosolic angle *θ*_*c*_. (b) ER and LD-tensions as functions of the cytosolic angle *θ*_*c*_. LD-monolayers have almost equal tensions and ER-tension is negligible during budding. (c) Almost identical *A*_*l*_’s in the two LD-monolayers as a function of the cytosolic angle *θ*_*c*_. The cytosolic angle *θ*_*c*_ varies almost linearly with the rescaled lipid number asymmetry Δ.

Similar to the LD transition simulations, we computed the ER and LD-tensions for the cytosolic LD budding. Unlike previous experiments that rely on tension imbalance between the two LD-monolayers to explain LD directional budding ^15^, we calculated equal tensions of about 4 *mN/m* in the two LD-monolayers, see Fig. 4b. Interestingly, the ER-bilayer remained almost tensionless during the LD budding from an ER-embedded spherical LD with *θ*_*c*_ = 92^*°*^ to an almost fully-emerged LD with *θ*_*c*_ = 163^*°*^.

Finding almost identical membrane tensions in the cytosolic and luminal LD-monolayers also implies approximately equal *A*_*l*_ values in the two LD-monolayers throughout LD budding. We verified this by computing *A*_*l*_ values for the cytosolic and luminal LD-monolayers as seen in the top panel of Fig. 4c. We obtained constant and almost identical *A*_*l*_ values in the two LD-monolayers of the emerging spherical LDs. The area of the cytosolic monolayer grows when adding more lipids. The area-per-lipid, however, remains almost constant, presumably with small fluctuations required for budding. Therefore, our results indicate that LD budding does not essentially require a significant tension imbalance between the cytosolic and luminal LD-monolayers. The bottom panel of Fig. 4c displays *θ*_*c*_ as a function of Δ for the emerging LDs in Fig. 4a. An almost linear growth is observed in *θ*_*c*_ upon increasing Δ.

### Free energy landscape of LD transition

The underlying mechanism of LD transition can be understood by identifying the free-energy contributions from bending the LD-monolayers, from ER and LD-tensions, and from the line tension. We calculated the free energy using a mean field approach, typically applicable to membranes where large bending and tension energies are minimally affected by thermal fluctuations as observed recently for membrane interactions with aqueous droplets ^57,59^. Note that the ER-bilayer, as a flat membrane with no spontaneous curvature, does not contribute to the bending energy. Figure 5a shows the total free energy and its different components for different symmetrical LDs, shown in Figure 2b, as a function of *θ* .

**Fig. 5.**
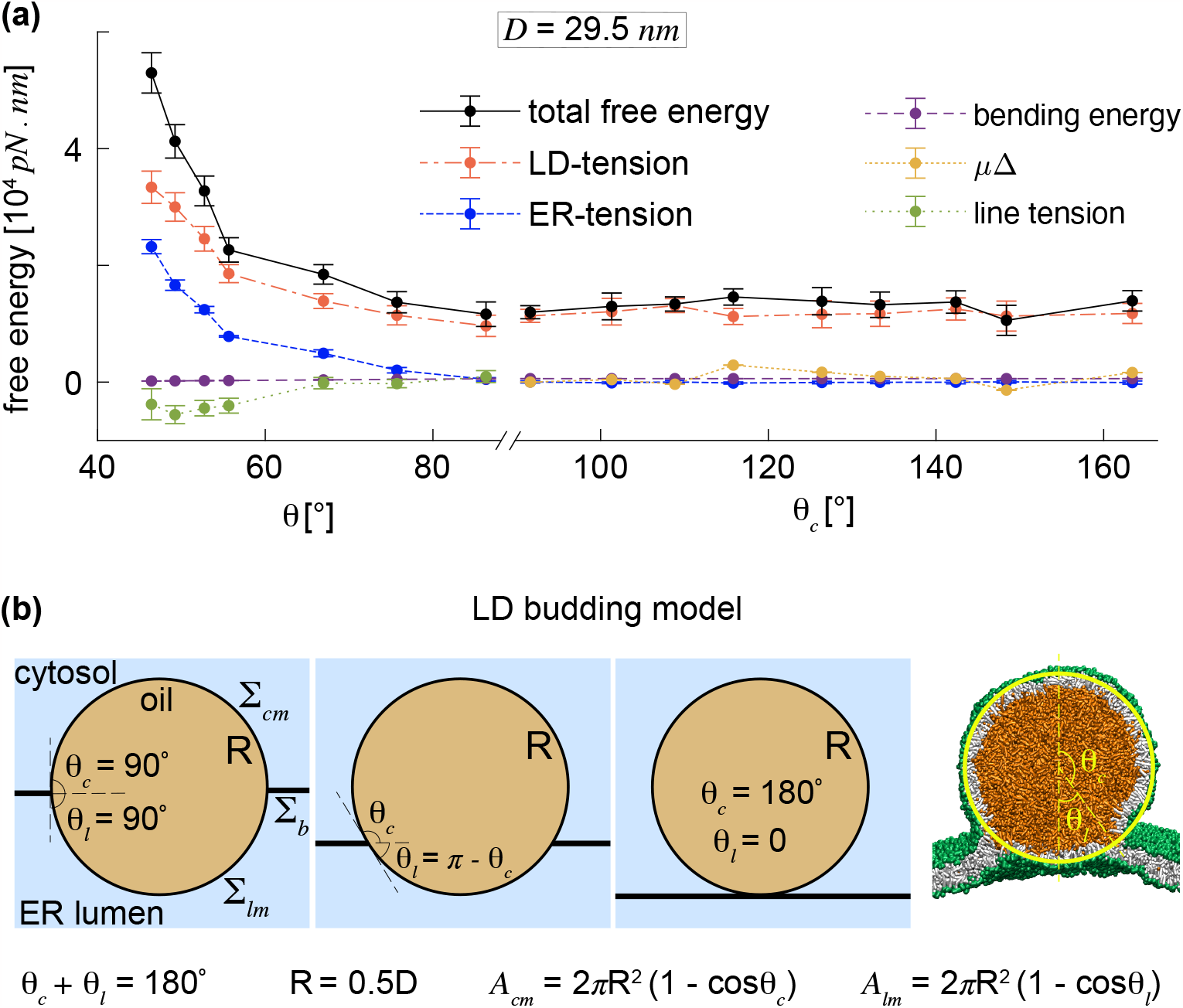
Free energy landscape of LD transition and budding. (a) Total free energy of an LD of size *D* = 29.5 *nm* as the sum of its components as a function of *θ* during LD transition and *θ*_*c*_ during LD budding. LD budding is energetically neutral as seen from the term *µ*Δ. (b) LD budding model used to describe the budding process of a spherical LD to the cytosol.

During the transition from an LD lens to an almost spherical LD with *θ* = 86^*°*^, the free energy rapidly dropped from about 5 *×* 10^4^ *pN · nm* to about 1 *×* 10^4^ *pN · nm* mainly due to the decreasing ER and LD-tensions, which compensated for the small increase in the line energy. In this regard, the LD transition from LD lenses to spherical LDs is similar to the formation of spherical oil droplets in water in the absence of lipid membranes. The latter one is also driven by the surface tension at the oil-water interface. This is reflected in the bending energy curve in Fig. 5a which barely contributes to the total free energy of the LD and thus to the LD transition. Overall the free energy of the LD transition indicates the negligible contribution of membrane bending and line tension even for the nano-sized LDs considered here. This is particularly important as the line tensions are generally expected to contribute more significantly at the sub-micron scale.

### Free energy landscape of LD budding

LD budding by creating an LD lipid number asymmetry, via lipid exchange at constant *N*_*lip*_, can be described by a simple budding model. During budding, the area of the cytosolic LD-monolayer increases while that of the luminal monolayer decreases upon exchanging lipids between LD-monolayers. Our simulations demonstrated that this process proceeds in such a way that the *A*_*l*_ values in LD-monolayers and thereby LD-tensions do not change during budding. This suggests that the initial symmetrical spherical LD can bud by retaining its spherical shape as a constant sphere of size *R* = 0.5*D*, see Fig. 5b. As a result, the cytosolic and luminal LD-monolayers can be represented by two spherical caps of this asymmetrical sphere (with respect to the ER-bilayer), each with their monolayer angles, *θ*_*c*_ and *θ*_*l*_ = *π* −*θ*_*c*_, respectively. The initial symmetrical sphere is characterized by *θ*_*l*_ = *θ*_*c*_ = *π/*2. The areas of the cytosolic and luminal LD-monolayers are given by the areas of the spherical caps, *A*_*jm*_ = 2*πR*^2^(1 − cos *θ* _*j*_ ) ( *j* = *c, l*).

We learned from free energy calculations of LD transition that energetic contributions of the line tensions are negligible. Therefore, we simplified our calculations by ignoring the line tension for LD budding. The constant bending energy, 8*πκ*_*m*_, of a spherical LD throughout budding does not contribute to the variation of the free energy. Here, *κ*_*m*_ is the bending stiffness of the LD-monolayers. Moreover, energetic contribution of membrane bending was also demonstrated to be negligible during LD transition. Hence, the free energy of LD budding is composed of the stretching energies of the ER and LD membranes as well as the free energy associated with the lipid exchange from the luminal to the cytosolic LD-monolayer. For the spherical LD of radius *R* connected to an ER bilayer of width *L*, the free energy is given by:

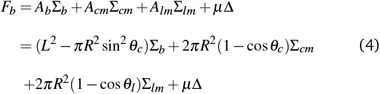

where Δ is given by Δ = *aθ*_*c*_ + *b* (fitted to the data in the bottom panel of Fig. 4c) and *µ* is the free energy conjugate of Δ. By setting *θ*_*l*_ = *π* − *θ*_*c*_ and ∑_*lm*_ = ∑_*cm*_, as obtained from simulations (Fig. 4b), and minimizing the resulting free energy with respect to *θ*_*c*_ we arrive at:

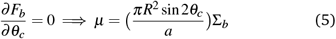

However, the ER-tension, ∑_*b*_, was demonstrated to be negligible for spherical LDs formed by LD transition (Fig. 2c) and remain negligible throughout LD budding (black dots in Fig. 4b). The equation 5 thus implies that *µ*, and thereby the free energy term *µ*Δ, are negligible. Consequently, LD budding by lipid exchange from the luminal to the cytosolic LD-monolayer, at constant total number of lipids, is energetically neutral.

Figure 5a shows the free energy contributions of membrane tensions, membrane bending, as well as the term *µ*Δ during LD budding for *θ*_*c*_ *>* 90^*°*^. The free energy *µ*Δ, associated to the lipid exchange is plotted with the actual values of *µ* calculated from Eq. 5, see the orange line in Fig. 5a. The free energy *µ*Δ is negligible compared to the other energetic contributions.

### Membrane tensions

The LD-tensions of about 4 *mN/m* that we found here for LDs of about 30 *nm* in diameter are very close to the LD tensions of ≈ 2 and ≈ 4 *mN/m* experimentally measured for LDs from mammalian and Drosophila cells ^14^. Comparing two different LD sizes, *D* = 27.7 and 29.5 *nm*, we observed almost identical ER and LD-tensions for both LDs, see supplementary Fig. S5. To examine the size-dependent behavior of LD-tensions, we computed these tensions for a very small LD of size *D* = 20.3 *nm*, and a larger one with *D* = 33.2 *nm* spontaneously formed from initial slabs with the same method as described earlier. Figure 6a shows ER and LD-tensions of these almost spherical LDs as a function of *D*. The LD-tensions were observed to slightly reduce with *D* for small LDs considered here. A more detailed analysis of size-dependent behavior of LD-tensions requires simulations of larger LDs, which will be accessible with future high performance computing.

**Fig. 6.**
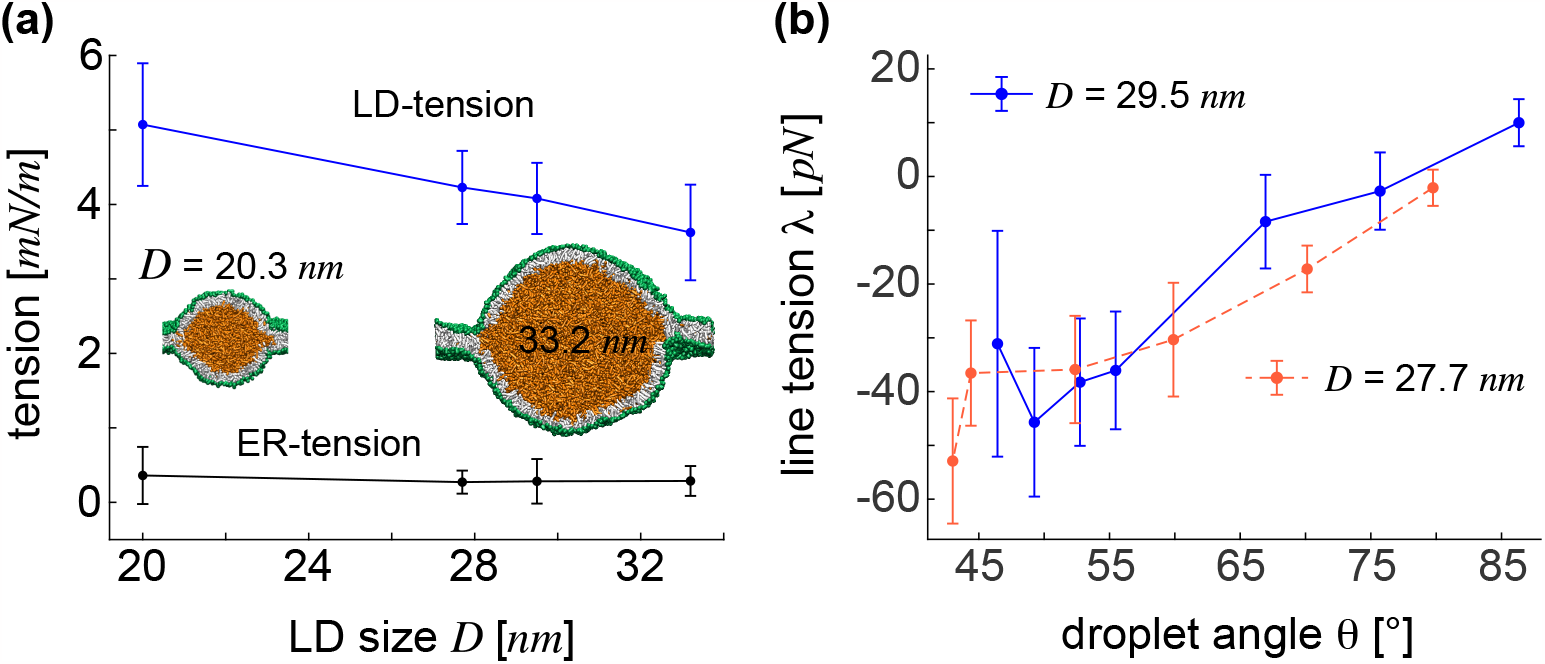
Size-dependent ER and LD tensions. (a) Size-dependent membrane tensions in four nearly spherical LDs of different sizes *D* = 20.3, 27.7, 29.5, and 33.2 *nm*. LD tension decreases with the LD size *D*. (b) Line tension along the LD contact line at the interface between neutral lipids, cytosolic water, and luminal water for two LDs of sizes *D* = 27.7 and 29.5 *nm*. Negative line tensions for small LD angles approach zero for spherical LDs.

The smallest LD of size *D* = 20.3 *nm*, we observed here, confirms our hypothesis on LD formation by size-independent transition process. These small spherical LDs have not been expected to form due to the dominant role of membrane bending ^11^ which is shown to be negligible in our simulations even in the limit of such a small LD size. According to our results, it is the size-independent tension balance that governs formation of different LD shapes through a continuous transition, as predicted earlier by the formation of multi-spherical LDs in Fig. S3.

We also computed the tension along the contact line of the interior oil with cytosolic and luminal aqueous solutions, see the Supplementary text for more details. Figure 6b displays the line tensions computed for two LDs of sizes *D* = 27.7 and 29.5 *nm*. Large negative line tensions appear to increase with lens angle *θ* amounting to a final negligible line tension for spherical LDs.

## Conclusion

We conducted extensive simulations of a coarse-grained model to gain insight into the mechanisms behind LD formation and budding with a primary focus on the mechanistic aspects of these processes. We began by identifying the relevant LD parameters, via spontaneous formation of LDs from slabs, which paved the way for our hypothesis that symmetrical LDs can form through a tension-induced continuous transition. We simulated several nano-sized LDs of different sizes and obtained various LD shapes for each LD size. The transition between LD lenses and spherical LDs appears to be continuous resembling a second-order phase transition. For small *A*_*lp*_ values together with single spherical LDs, we also identified the presence of smaller multi-spherical LDs, even though they are unstable (Fig. S3), suggesting the stability of nano-sized spherical LDs. These findings align with the principles governing oil droplet formation in water in the absence of lipids. This underscores the dominant role of membrane tensions, which are a common factor in both scenarios, and suggests the likely negligible contribution of membrane bending and line tensions in LD formation.

LD formation and budding are known to be regulated by membrane tensions ^11,14,28^. Both the formation of symmetrical ER-embedded LDs and the consequent budding of asymmetric LDs depend on ER and LD-tensions. It is shown experimentally that these tensions are proportional to lipid densities of the ER and LD membranes ^18^. The interplay between the ER and LD-tensions has been also shown to determine the LD shape and its transition to fully budded LDs that emerge to the cytosol ^11,14^. Previous studies have described LD transition and budding in terms of a discontinuous second-order transition which depends on the LD size, membrane tensions, and membrane bending ^11,28^.

Symmetrical LDs have been suggested to form through a size-dependent process, primarily controlled by membrane bending for small LD sizes and by membrane tensions for larger LDs ^11^. Consequently, symmetrical spherical LDs only formed for very large LD sizes while small LD sizes resulted in wide LD lenses with small lens angles. In our simulations, however, we observed spherical LDs even for small nano-sized LDs. This observation was further supported by calculating negligible energetic contributions of membrane bending and line tensions, in comparison to the contributions from membrane tensions.

We also demonstrated that the symmetrical LDs can form through a continuous first-order transition at fixed LD size, governed by the mechanical force balance between LD and ER tensions. The slight deviation from this force balance gives rise to a tension difference (Δ∑ *>* 0), which pulls the ER bilayer and triggers the continuous transition from tense LD lenses to spherical LDs with negligible ER-tensions. Unlike in previous studies where Δ∑ increased with *θ* ^11^, our calculations exhibit a reduction in Δ∑, and thus the pulling force on the ER-bilayer, as we approach the spherical LDs. This makes sense as the initial strong tendency of LD transition is expected to decrease for more spherical LDs, with increasing values of *θ* .

Similar to LD formation, the subsequent LD budding and directional emergence toward the cytosol has been also linked to the interplay of ER and LD-tensions and the LD size ^11^. Over a wide range of LD sizes, LD budding is believed to occur through a discontinuous first-order transition that transforms symmetric LD lenses with large ER-tension to asymmetric fully-budded LDs with small ER-tensions. By contrast, we demonstrated here via calculating free energies, that LD budding by lipid exchange between the LD-monolayers is energetically neutral. Unlike LD formation, which is primarily regulated by membrane tensions, LD budding is mainly driven by this lipid exchange, independent from membrane tensions and the LD size. The modest energy required for budding is presumably provided by proteins that act as the driving agents.

Energetically neutral budding by inter-monolayer lipid exchange is primarily due to the identical mechanical tensions in the luminal and cytosolic LD-monolayers. While the number of lipids and the area of the luminal and cytosolic LD-monolayers change by lipid exchange, their lipid densities or areas-per-lipid remain constant. This is the key to constant identical tensions in LD-monolayers and the consequent neutral budding (see Eq. 4) which makes LD budding a tension-independent process.

LD formation through size-independent continuous transitions of LD lenses and energetically neutral LD budding that we have discovered, carry significant implications. Prior understanding suggested that fully-budded asymmetric LDs were the stable structures for sufficiently small ER-tensions across a wide spectrum of LD sizes ^11^. This understanding made it challenging to explain the stability of experimentally-observed symmetric LDs ^12,14^, particularly those with more spherical shapes. However, our new findings, which reveal a continuous size-independent LD transition driven by the reduction in free energy, followed by energetically-neutral budding, provide a clear explanation for the stability of these close-to-spherical LDs.

The role of protein machinery in LD budding remains a contentious topic in the field ^1^. While certain studies have asserted that proteins actively participate in LD budding, others have considered proteins as passive observers with no significant role ^1,31,32^. The energetically-neutral nature of LD budding that we have uncovered, based on inter-monolayer lipid exchange, provides new insight into the role of proteins in LD budding. On one hand, we demonstrate that LD budding occurs neutrally, with no remarkable energy input required. On the other hand, any energetically-neutral process demands a slight energy input to take place in a quasi-equilibrium manner. Our findings thus suggest that proteins are indeed essential for LD budding, but their energetic contributions are modest. This perspective bridges the gap between the existing opposing views on the role of proteins in LD budding.

Despite the coarse-grained nature of our molecular simulations with soft potentials, we computed very accurate ER and LD-tensions. LD-tensions we computed here for nearly spherical ER-embedded LDs of different sizes, between 3.5 and 5 *mN/m* as seen in Fig. 6a, are surprisingly close to the LD-tensions of about 3.5 *±* 1.5 and 2 *±* 0.5 *mN/m* measured experimentally for LDs from mammalian and Drosophila cells, respectively ^14^.

The negative line tensions, we calculated here for wide LD lenses, increase to approach positive values for almost spherical ER-embedded LDs, see Fig. 6b. These tensions presumably remain positive during the entire budding of almost spherical LDs. To the best of our knowledge, tensions of contact lines between lipid droplets and lipid bilayers have not been measured experimentally. Whether the negative line tension, reported here for wide LD lenses, is an artefact of the DPD force field or they can actually exist for LDs in the ER awaits further computational and experimental studies. In particular, such future studies could involve other computational modes, such as the Martini force field ^64^ among other available coarse-grained models.

## Author Contributions

A.H.B. designed research; F.K., B.R., and A.H.B. performed research; F.K., A.B., R.G., B.R., and A.H.B. analyzed the data; A.H.B. wrote the paper with substantial contributions from F.K., R.G., and B.R.

## Acknowledgements

A.H.B. acknowledges support from the Max Planck Society within the framework of Max Planck Partner Group, and from the European Molecular Biology Organization, grant EMBO IG 5032. B.R. acknowledges support from the National Science Centre, Poland, grant no 2020/39/B/NZ1/00377.

## Supplementary Information

### Supplementary Text

#### S1. Assembling initial slabs

We started by assembling initial slabs, wherein a given volume *V* of oil is covered by two flat lipid monolayers from below and above with their hydrophobic chains facing the oil, see Fig. 1a and supplementary Fig. S1b. Initial monolayers were assembled by placing different number of lipids *N*_*lip*_, equally divided between two monolayers of fixed width *L* in a cubic simulation box with periodic boundary conditions. Each slab with a given number of lipids *N*_*lip*_ has then a fixed projected area-per-lipid *A*_*lp*_ = 2*A*_*s*_*/N*_*lip*_, defined as the average area-per-lipid projected into the horizontal plane.

#### S2. Shape parameters of LDs

We first computed the thickness of the ER-bilayer *l*_*b*_ = 4.08 *nm* as the distance between the peaks of the density of lipid head groups of a flat bilayer, see supplementary Fig. S1a. The monolayer thickness was then taken to be *l*_*m*_ = 0.5*l*_*b*_ = 2.04 *nm*. We assumed that LD-monolayers of all LDs, obtained from DPD simulations, have the shape of spherical caps to minimize membrane bending cost for a given membrane surface area. To find the shape of symmetrical LD lenses and spherical LDs, we fitted spherical caps to the simulation results and computed their shape parameters according to Figure 2a. Complete spheres of size *R* = 0.5*D* were fitted to the asymmetric budding LDs. The contact point of the ER-bilayer with these spheres were then used to find the LD angles as shown in the right panel of Fig. 5b. The spherical caps of symmetrical LDs were parametrized by the radius of the sphere and the Cartesian coordinates *x*_*s*_, *y*_*s*_, and *z*_*s*_ of its center. The sum *S* of the mean square distances of all beads *i* of LD lipids from the surface of the cap

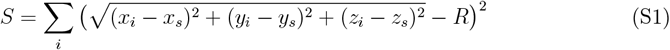

was then minimized by setting the derivatives

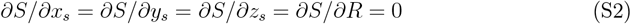

leading to a system of four equations from which the radius *R* of the cap and the Cartesian coordinates of the cap center were obtained. The contact angle was then computed from the fitted cap. In this way, we fitted spherical caps to LD-monolayers and obtained the radii and lens angles of the luminal and cytosolic LD-monolayers for different LD shapes.

#### S3. Membrane tension and bending stiffness

To find membrane properties such as membrane thickness and bending stiffness of bilayers and monolayers, we simulated small flat bilayers and small slabs. The slabs were simulated in a cubic box of fixed width 25.6 *nm* and initial height 25.6 *nm* with *N*_*oil*_ = 10, 363 oil molecules surrounded by 35, 852 water beads forming slabs of size *D* = 24.2 *nm*, see Fig. 1a. The number of lipids *N*_*lip*_, equally partitioned between the two slab monolayers, was varied to generate slabs with different *A*_*lp*_, see supplementary Table S2. Flat bilayers were also simulated in cubic boxes of the same size with the same *N*_*lip*_ and thus the same *A*_*lp*_ as the slabs, see the snapshot in supplementary Fig. S2c and the data in supplementary Table S2.

Both slabs and flat bilayers were first equilibrated for 2 *µs* in an NPT ensemble using a Berendsen barostat in the vertical direction *z*, see supplementary Fig. S2a,c, adjusting the pressure at 23.7 *k*_*B*_*T/d*^3^ = 190.4 *×* 10^9^ *mN/m*^2^. During NPT simulations the height of the simulation boxes changed to adjust the pressure while their widths remained constant at 25.6 *nm*. We note that the density of the simulation box, including both water and oil, slightly varies during NPT simulations but the water density remains at 5.86*/nm*^3^ as shown in the density plots in the bottom panels of Fig. S1. Membrane tensions were then computed in following 22 *µs*-long runs in an NVT ensemble with fixed simulation boxes at room temperature. We used the integral

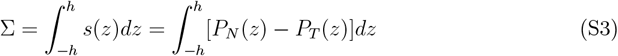

of the stress profile *s*(*z*) = *P*_*N*_ (*z*) − *P*_*T*_ (*z*) to find both monolayer and bilayer tensions, ∑_*m*_ and ∑_*b*_, respectively. Here, *P*_*N*_ and *P*_*T*_ denote normal and tangential pressure components, computed across an almost planar patch of a bilayer or a monolayer membrane in the *z* direction perpendicular to the membrane surface. 1,2 We first calculated membrane tensions inside small slabs of size *D* = 24.2 *nm* to demonstrate that monolayer tensions increase linearly with *A*_*lp*_. The Cartesian coordinate system was placed in the center of the slabs and the stress profile was computed inside a box of width 13 *nm* and height 2*h* = 20 *nm*, see supplementary Fig. S2a. The monolayer tensions were then calculated as

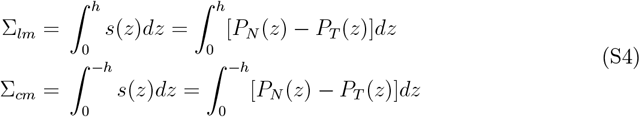

The monolayer tension ∑_*m*_ = 0.5(∑_*lm*_ + ∑_*cm*_) was taken to be the average of almost equal tensions in the outer (cytosolic) and inner (luminal) monolayers. Supplementary Figure S2b shows the stress profiles of some of these small slabs with corresponding ∑_*m*_’s and *A*_*lp*_’s whose values are listed in supplementary Table S2.

We computed the monolayer tension ∑_*m*_ in the initial flat monolayers of three small slabs of size *D* = 24.2 *nm* and area *A*_*s*_ = 655.4 *nm*^2^, with different *A*_*lp*_’s, see Fig. 1b. The slab area was chosen to be smaller than *A*_*sm*_ = 1115.3 *nm*^2^ for *L* = 25.6 *nm* to prevent the slab-to-lens transition. The monolayer tension increased from 0.4 *mN/m* for *A*_*lp*_ = 0.72 *nm*^2^ to 10.5 *mN/m* for *A*_*lp*_ = 0.82 *nm*^2^ corresponding to monolayers with *N*_*lip*_ = 1800 and *N*_*lip*_ = 1600 lipids, respectively, see supplementary Fig. S2b and Table S2. We thus verified that lipid membranes with larger *A*_*lp*_’s experience larger membrane tensions.

We then used equation (S3) to obtain stress profiles of flat bilayers in a box of width 13 *nm* and initial height *h* = 5 *nm*, see supplementary Fig. S2c. Supplementary Figure S2d shows the resulting stress profiles across two flat bilayers. For both bilayers and monolayers (slabs), membrane tensions and areas-per-lipid were averaged over 400 simulation frames. These simulations were performed for 20 *µs* in an NVT ensemble during which 400 equally distanced frames were picked to compute the average ∑’s and *A*_*lp*_’s as listed in supplementary Table S2. These tensions, plotted against the corresponding *A*_*lp*_ values by blue and red dots in supplementary Fig. S2e, were then used to compute the bending rigidities of the ER-bilayer and LD-monolayers.

We fitted a line to the data of ∑_*m*_ versus *A*_*lp*_, and found 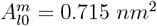 corresponding to zero tension ∑_*m*_ = 0. The area extension elastic modulus was then obtained as the product 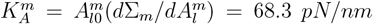, where 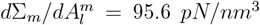.^3^ The bending rigidity of the monolayer was then found as 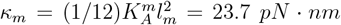, where *l*_*m*_ = 2.04 *nm* is the monolayer thickness i.e. half the bilayer thickness *l*_*m*_ = 0.5*l*_*b*_. A similar procedure was used to find the bending rigidity of the ER-bilayer as:

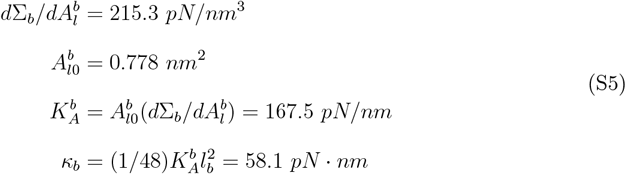

In this way, we found the bending rigidities, *κ*_*b*_ = 58.1 *pN · nm* and *κ*_*m*_ = 23.7 *pN · nm*, of the ER-bilayer and LD-monolayers, respectively.

#### S4. LD parameters

Spontaneous formation of LDs from initial oil slabs were simulated in a fixed simulation box of size (48 *×* 48 *×* 72) *nm*^3^ composed of *N*_*lip*_ lipids placed in two monolayers exposed to water from above and below with a fixed total of 972000 beads. The number of lipids was then varied, as listed in supplementary Table S3, to change the projected area-per-lipid *A*_*lp*_ for different LD sizes *D* = 29.5, 31, and 33.2 *nm* as seen in the morphological diagram in Fig. 1c. Spontaneous formation of different LD structures occurred in 5 *µs* in an NPT ensemble at constant pressure 190.4 *×* 10^9^ *mN/m*^2^, using a Berendsen barostat in the vertical direction *z* perpendicular to the slab monolayers. The initial slabs transformed to a variety of LD structures as shown in Fig. 1c.

#### S5. The transition of LD lenses to spherical LDs

We performed the transition simulations starting from an almost spherical LD of size *D* = 29.5 *nm* with the largest lens angle *θ* = 86^*°*^, spontaneously formed from an initial slab with *N*_*oil*_ = 21, 000 and *N*_*lip*_ = 6960. We then increased the width *L* of the simulation box in an NPT ensemble with a barostat in the vertical direction *z* perpendicular to the ER-bilayer for a constant number of lipids and water beads.

We increased the width of the simulation box at a rate 0.4 *pm/ns* resulting in a quasi-equilibrium transition with intermediate equilibrium LD lenses.^4^ The procedure was then repeated successively to find different lenses shown in Fig. 2b. After reaching the target box width for each lens angle, we continued the simulation for 8 *µs*, this time in the NVT ensemble, the last 6 *µs* of which was used to compute membrane tensions by averaging over 120 frames, and *A*_*l*_ values, and LD shape parameters by averaging over 12 frames. LD-tensions were computed, using the equations (S4), in a box of width 8 *nm* and height 48 *nm*, which contained almost flat monolayer patches as shown in supplementary Fig. S4. ER-tensions were computed using equation (S3) inside a box of size (8 *×* 48 *×* 48) *nm*^3^, see supplementary Fig. S4. The same boxes were used to compute *A*_*l*_ values for both the ER-bilayers and LD-monolayers. Supplementary Table S4 lists the LD parameters for the resulting seven LDs shown in Fig. 2b. To compare different LD sizes, we also performed the transition simulations for another LD of size *D* = 27.7 *nm* with *N*_*oil*_ = 16800 and *N*_*lip*_ = 6800, the parameters of which are presented in supplementary Table S5. See the supplementary text for details on line tension calculation during LD transition.

#### S6. Line tensions in symmetrical LDs

The free energy of a symmetrical LD with the surrounding ER-bilayer is given by

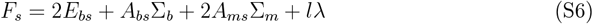

Here, *E*_*bs*_ = 2*κ*_*m*_*A*_*ms*_*/R*^2^ is the bending energy of the two symmetrical LD-monolayers, ∑_*b*_ and ∑_*m*_ are the ER and LD-tensions, and *λ* is the line tension acting along the perimeter *l* = 2*πR*_*co*_ of the contact circle with radius *R*_*co*_. *A*_*bs*_ = *L*^2^ − *πR*^2^ sin^2^ *θ* is the area of the surrounding ER-bilayer of width *L, A*_*ms*_ = 2*πR*^2^(1− cos *θ*) is the surface area of the spherical monolayers with identical radii *R*.

Minimizing the Lagrangian *ℒ*_*s*_ = *F*_*s*_ − *γV*_*s*_ with respect to *R* and *θ* using a Lagrange multiplier *γ* to fulfil a constraint on fixed LD size *V*_*s*_ = *πD*^3^*/*6, leads to a system of three equations

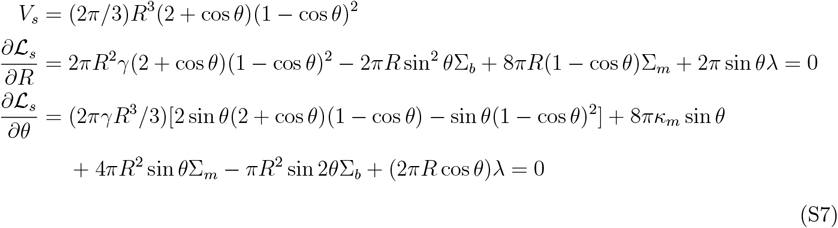

with four variables *R, θ, λ, γ*. To solve this underdetermined system, we substituted the values of *R*, ∑_*b*_, and ∑_*m*_, obtained from the DPD simulations, into the system and solved equations (S7) for the remaining three unknowns *λ, θ*, and *γ*. The results for two LD sizes are listed in Tables S4 and S5.

#### S7. Membrane elasticity model for axisymmetric LDs

To determine the minimum energy shapes of symmetric lenses, we adapted a computational approach used in^5^ based on the elasticity theory of fluid membranes. Here, the bottom monolayer of the lens is described as a smooth surface in terms of two parameters, the angle *γ* of rotation around the axis of symmetry, and the arc length *s* along longitudes. This surface then has Cartesian coordinates *X* = *R*(*s*) cos *γ, Y* = *R*(*s*) sin *γ* and *Z* = *Z*(*s*), where 0 ≤ *γ* ≤ 2*π* and 0 ≤ *s* ≤ *s*_1_ and *R* is the distance from the *Z*-axis, which is the axis of symmetry. We introduce a tangent angle Ψ such that *dR/ds* = cos Ψ(*s*) and *dZ/ds* = sin Ψ(*s*). In this parametrization, the volume of the lens is

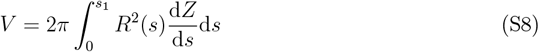

and the total area of the two monolayers enclosing the lens is

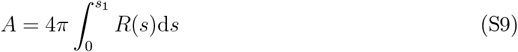

The lower limit of integration, *s* = 0, corresponds to the “south pole” of the lens at which Ψ(*s* = 0) = 0, *R*(*s* = 0) = 0 and *Z*(*s* = 0) = 0. With these boundary conditions we have 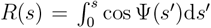 and 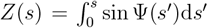. The upper limit of integration, *s* = *s*_1_, corresponds to the lens rim, where Ψ(*s*_1_) = 0 and *R*(*s*_1_) = *R*_1_ to connect the lens surface smoothly to a flat bilayer at a distance *R*_1_ from the axis of symmetry. The circumference of this rim is

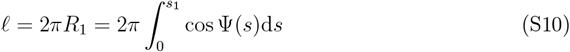

The energy of the system *E* = *E*_b,m_ + *E*_b,b_ + *E*_s,m_ + *E*_s,b_ + *E*_l_ comprises five terms: the bending energy *E*_b,m_ of the two monolayers, the bending energy of the flat bilayer, *E*_b,b_ = 0, the surface energy *E*_s,m_ of the monolayers, the surface energy *E*_s,b_ of the bilayer, and the line energy *E*_l_ of the lens rim. Since the principal curvatures of the lens surface are given by *C*_1_ = *d*Ψ*/ds* and *C*_2_ = sin Ψ*/R* in the arc-length parametrization,^6,7^ the bending energy of the two monolayers is

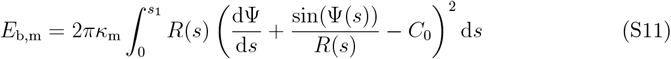

where *κ*_m_ and *C*_0_ are the bending rigidity modulus and the spontaneous curvature of the monolayers, respectively. The surface energy of the monolayers *E*_s,m_ = *A*∑_s_, where ∑_s_ denotes the surface tension of the monolayers, and the lens surface area *A* is given by equation (S9). The surface energy of the bilayer 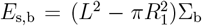, where ∑_b_ denotes the surface tension of the bilayer, and *L* is the lateral size of the system. The energy term *L*^2^∑_b_ does not depend on the shape of the lens and can be omitted in numerical calculations. Finally, the line energy *E*_l_ = *λf*, = 2*πR*_1_*λ*, where *λ* denotes the line tension. It is convenient to use dimensionless variables in numerical calculations. Here we introduce *τ* = *s/s*_1_ with *τ* ∈ [0, 1], *ψ*(*τ* ) = Ψ(*s*), *r*(*τ* ) = *R*(*s*)*/s*_1_ and *z*(*τ* ) = *Z*(*s*)*/s*_1_. Then, using equation (S8), *s*_1_ can be directly related to the volume *V* of the lens

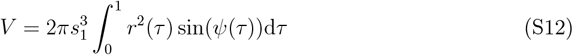

In addition, with this choice of dimensionless variables, the total energy of the system can be written as *E* = *E*_b_ + *E*_m_ + *E*_l_ with

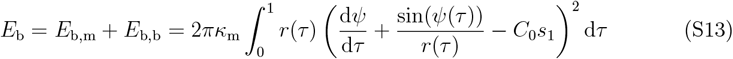

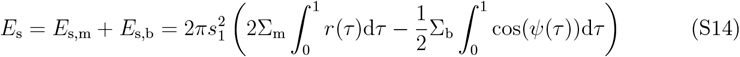

and

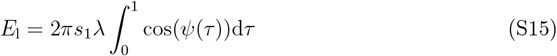

In equations (S13), (S14) and (S15), parameters *κ*_m_, *C*_0_, ∑_m_, ∑_b_, and *λ* characterize the mechanical properties of the system whereas *s*_1_ is determined by the volume *V* of the lens *via* equation (S12). Since *r*(*τ* ) and *z*(*τ* ) are given by *ψ*(*τ* ) *via*

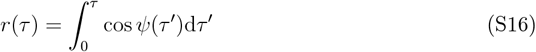

and

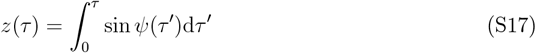

the shape of the lens is entirely determined by the function *ψ*(*τ* ), and the total energy *E* = *E*_b_ + *E*_m_ + *E*_l_ is a functional of *ψ*(*τ* ).

To minimize the total energy *E* with respect to the lens shape, as given by the function *ψ*(*τ* ), it is convenient to approximate *ψ*(*τ* ) by a Fourier series^8,9^

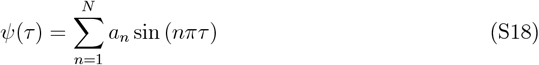

that fulfils the boundary conditions *ψ*(*τ* = 0) = 0 and *ψ*(*τ* = 1) = 0. Here, *N* is the number of Fourier amplitudes *a*_*n*_. The total energy *E* = *E*_b_ + *E*_m_ + *E*_l_, as given by equations (S13), (S14) and (S15), becomes now a function of *N* variables {*a*_*n*_}.

We minimized *E* with respect to the set of Fourier amplitudes {*a*_*n*_} using a simulated annealing method. We performed the numerical calculations with *N* = 100 Fourier modes as in.^8,9^ We assumed *C*_0_ = 0 whereas the values of parameters *κ*_m_, ∑_m_, ∑_b_, *λ* and *V* were taken from the optimal fits of the spherical cup model to the lens shapes obtained in the DPD simulations. The resulting membrane profiles *ψ* = *ψ*(*s*) are shown in Figure 3 by yellow curves juxtaposed on DPD simulation snapshots for different LD lenses.

#### S8. LD budding by lipid exchange

Starting from an almost spherical LD of size *D* = 29.5 *nm*, we created an asymmetric lipid number at constant *N*_*lip*_ (see Table S6) by exchanging lipids from the luminal to the cytosolic LD-monolayer. The resulting LDs with a given Δ were then equilibrated for 2 *µs* in the NPT ensemble at constant pressure 190.4 *×* 10^9^ *mN/m*^2^ using a Berendsen barostat in the vertical direction. The equilibration was followed by a 6 *µs*-long NVT simulation during the last 4 *µs* of which membrane properties were averaged over 80 simulation frames. Supplementary Table S6 lists the LD parameters for the resulting LDs shown in Fig. 4a with different cytosolic angles *θ*_*c*_.

#### S9. Size-dependent LD-tension

Figure 2c shows ER and LD-tensions as a function of *θ* for an LD of size *D* = 29.5 *nm*. To estimate finite size effect on the LD-tensions, we repeated the same calculations for a smaller LDs of size *D* = 20.3 *nm* and 27.7 *nm*. Although LD-tensions inside spherical LDs seem to be smaller for the larger LD, see supplementary Fig. S5, their values are within the errorbars of one another. Therefore, we simulated more LDs with different sizes, four of which were used to explore size-dependent behavior of LD-tensions as shown in Fig. 6a. Supplementary Table S7 lists LD parameters for these four LDs which spontaneously formed from initial slabs.

## Supplementary Figures

**Figure S1:**
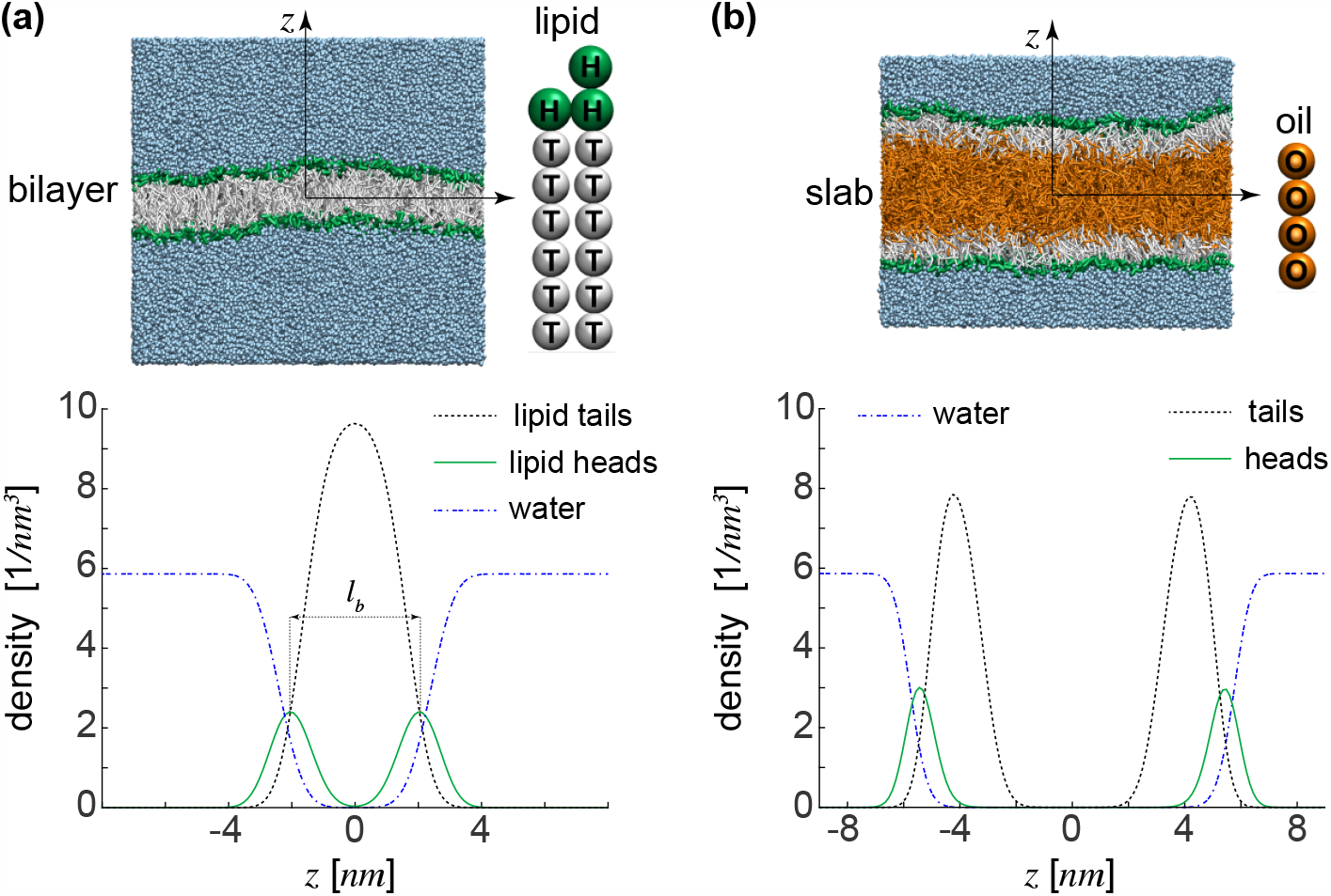
Coarse-grained membrane and neutral lipids with their densities: (a) A lipid bilayer with the corresponding density profiles of lipid heads, lipid tails, and water beads. The distance between the peaks of the lipid heads density defines the bilayer thickness *l*_*b*_. Also shown is the structure of a lipid with its head *H* and tail *T* beads. (b) The density profiles of lipid tails and heads, and water for a slab of size *D* = 24.2 *nm*. As expected water has a density of 3*/d*^3^ = 5.86*/nm*^3^. Also shown is an oil molecule with its oil beads *O*.

**Figure S2:**
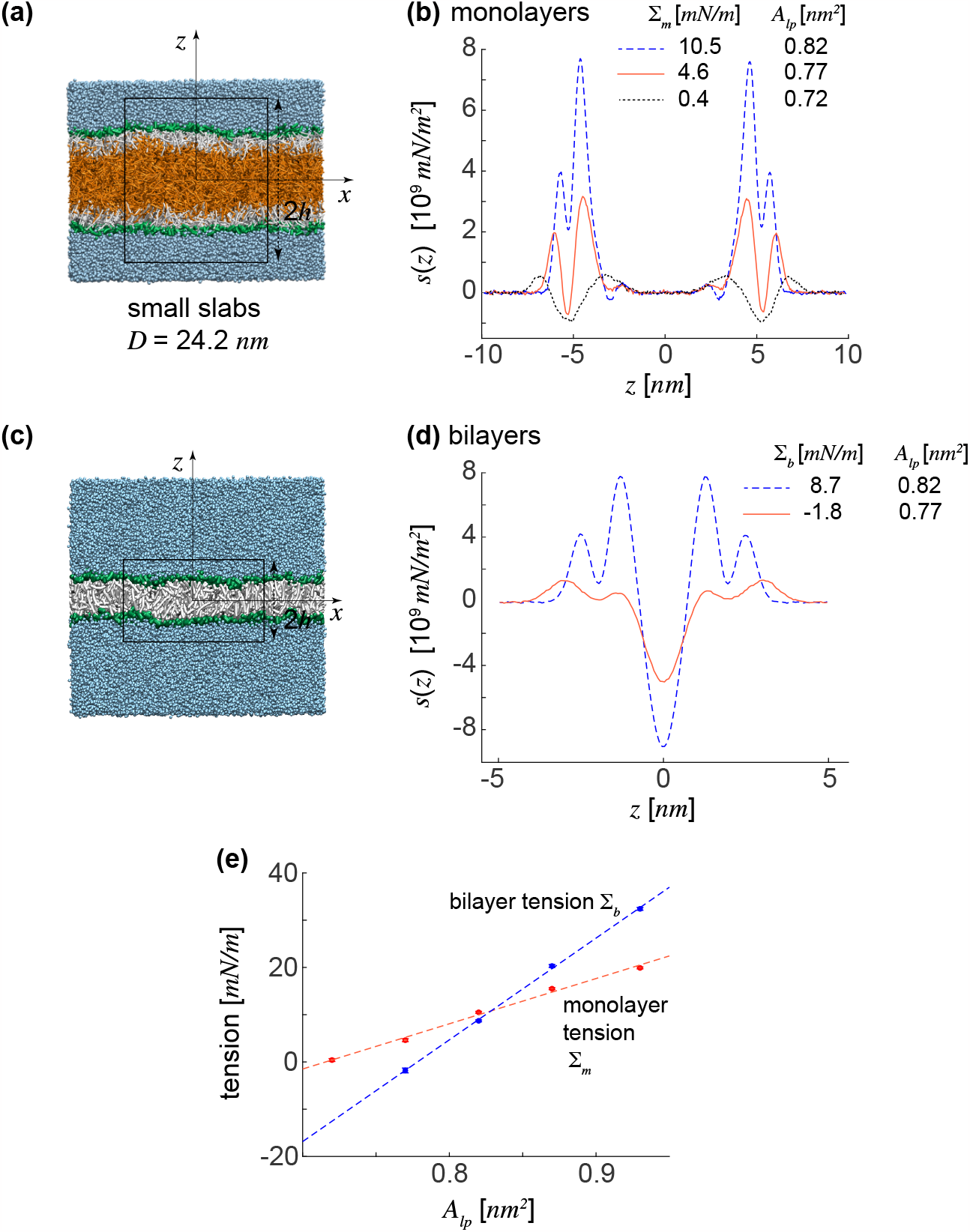
Finding membrane stiffness and thickness: (a) Small slabs of size *D* = 24.2 *nm* used to compute bending stiffness and membrane thickness of LD-monolayers with boxes of height 2*h* = 20 *nm* and width 13 *nm* for calculating stress profiles as shown in (b) for three slabs. (c) Small flat bilayers with boxes of height 2*h* = 10 *nm* and width 13 *nm* for calculating stress profiles in bilayers as shown in (d) for two bilayers. (e) Monolayer and bilayer tensions, ∑_*m*_ and ∑_*b*_, plotted against *A*_*lp*_ with red and blue dots. The dashed lines, fitted to the data, were used to calculate membrane stiffness.

**Figure S3:**
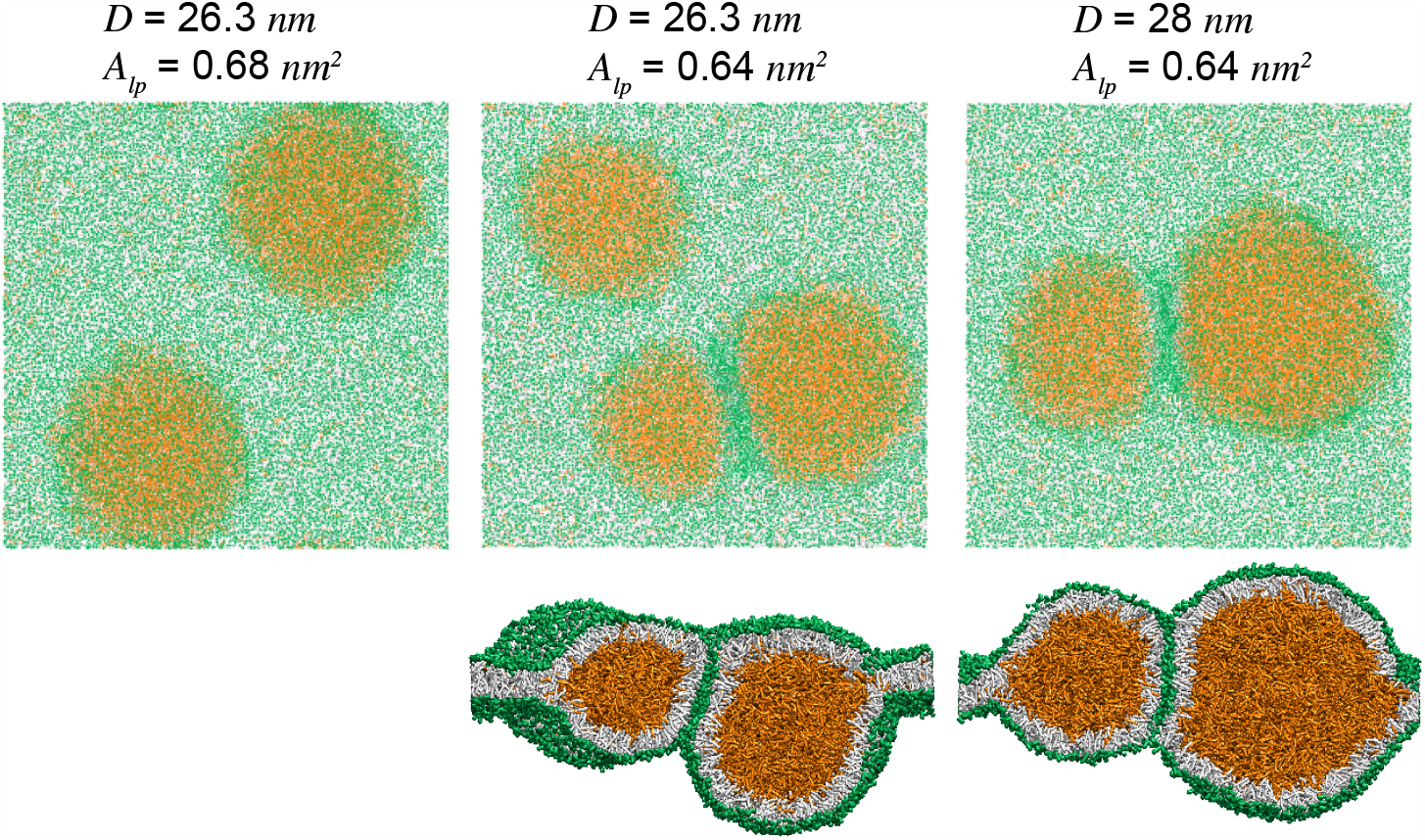
Multi-spherical LDs: In addition to single spherical LDs, multi-spherical LDs of smaller sizes form for sufficiently small area-per-lipids of initial slabs. Green and orange dots represent oil and lipid molecules, respectively. The bottom snapshots of the middle and right panels show the lateral cross sections of double LD structures.

**Figure S4:**
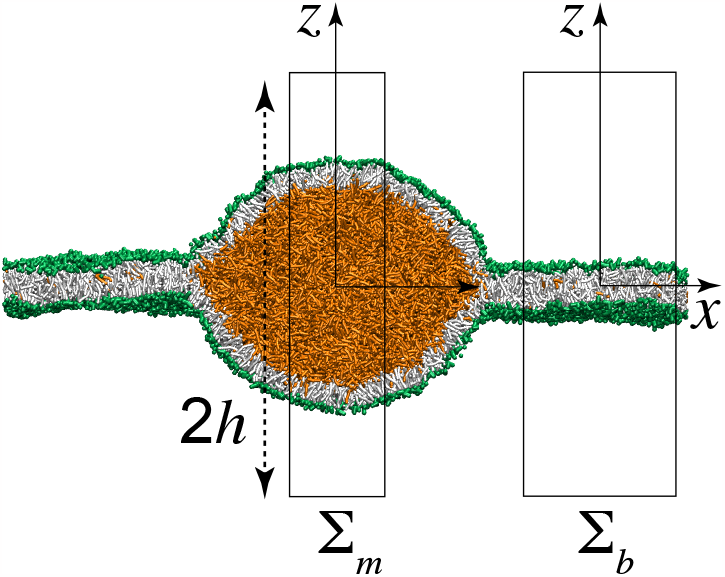
Stress calculation boxes: Boxes of width 5 and 8 *nm* and height 48 *nm* used to compute LD and ER-tensions, ∑_*m*_ and ∑_*b*_, and their areas-per-lipid.

**Figure S5:**
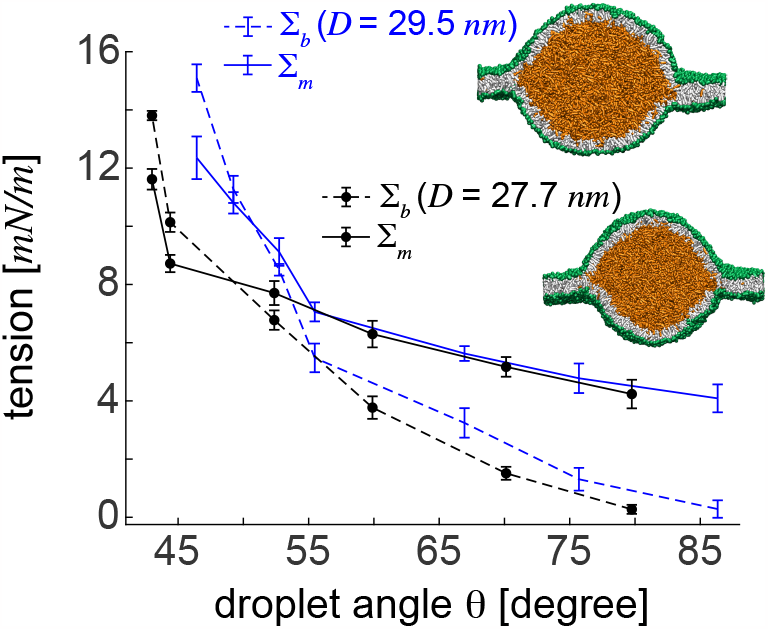
Size-dependent behaviour of ER and LD-tensions: ER and LD-tensions versus lens angle *θ* for two LDs of sizes *D* = 29.5 and *D* = 27.7 *nm*. LD-tensions of almost spherical LDs seem to be almost invariable for the two LD sizes.

## Supplementary Tables

**Table S1:** DPD Interaction force parameters *f*_*ij*_ [*k*_*B*_*T/d*]

| | $j = W$ | $j = H$ | $j = T$ | $j = O$ |
| --- | --- | --- | --- | --- |
| $i = W$ | 25 | 30 | 75 | 75 |
| $i = H$ | 30 | 30 | 50 | 50 |
| $i = T$ | 75 | 50 | 10 | 11 |
| $i = O$ | 75 | 50 | 11 | 10 |

The interaction force parameter *f*_*ij*_ between oil *O*, water *W*, lipid head *H*, and lipid tail *T* beads.

**Table S2:** Properties of small slabs and flat bilayers

| $N_{lip}$ | $A_l^m, A_l^b$ [$nm^2$] | $\Sigma_m$ [$mN/m$] | $\Sigma_b$ [$mN/m$] |
| --- | --- | --- | --- |
| 1800 | 0.72 | $0.4 \pm 0.6$ | — |
| 1700 | 0.77 | $4.6 \pm 0.3$ | $-1.8 \pm 0.5$ |
| 1600 | 0.82 | $10.5 \pm 0.2$ | $8.7 \pm 0.2$ |
| 1500 | 0.87 | $15.5 \pm 0.2$ | $20.3 \pm 0.3$ |
| 1400 | 0.93 | $19.9 \pm 0.2$ | $32.4 \pm 0.3$ |

Areas-per-lipid and tensions of flat bilayers and small slabs of size *D* = 24.2 *nm* (∑_*b*_ and ∑_*m*_ respectively) used to find the bending stiffness and membrane thickness of the ER-bilayer and LD-monolayers. All slabs contain the oil volume *V* = 4264.6 *nm*^3^ with slab area *A*_*s*_ = 655.4 *nm*^2^ and minimum slab area *A*_*sm*_ = 1115.3 *nm*^2^. The monolayer thickness is *l*_*m*_ = 2.04 *nm*.

**Table S3:** Properties of initial slabs used for identifying relevant LD parameters

| $N_{oil}$ | $D$ [nm] | $V$ [nm<sup>3</sup>] | $N_{lip}$ | $A_{lp}$ [nm<sup>2</sup>] | $A_{sm}$ [nm<sup>2</sup>] |
| --- | --- | --- | --- | --- | --- |
| 21000 | 29.5 | 8601.6 | 6800 | 0.68 | 2987.5 |
|  |  |  | 6400 | 0.72 |  |
|  |  |  | 6000 | 0.77 |  |
|  |  |  | 5600 | 0.822 |  |
| 24881 | 31 | 10192 | 7200 | 0.64 | 3058.8 |
|  |  |  | 6428 | 0.716 |  |
|  |  |  | 5625 | 0.819 |  |
| 31500 | 33.2 | 10902.4 | 6800 | 0.68 | 3169.7 |
|  |  |  | 6400 | 0.72 |  |
|  |  |  | 6000 | 0.77 |  |

Properties of slabs with different sizes used for spontaneous formation of LDs and identifying LD parameters. The data corresponds to slabs the resulting LDs of which are shown in Figure 1c. The slab area is *A*_*s*_ = 2304 *nm*^2^ and the monolayer thickness is *l*_*m*_ = 2.04 *nm* for all slabs.

**Table S4:** Properties of symmetrical LD lenses with size *D* = 29:5 *nm*

| $L$ [nm] | $R$ [nm] | $\theta$ [°] | $\Sigma_b$ [mN/m] | $\Sigma_m$ [mN/m] | $\lambda$ [pN] | $\gamma$ [10<sup>9</sup> mN/m<sup>2</sup>] |
| --- | --- | --- | --- | --- | --- | --- |
| 47.2 | $14.2 \pm 0.3$ | $86.3 \pm 0.1$ | $0.3 \pm 0.3$ | $4.1 \pm 0.5$ | $10 \pm 4.4$ | $-0.615$ |
| 48 | $15.9 \pm 0.3$ | $75.7 \pm 0.07$ | $1.3 \pm 0.4$ | $4.8 \pm 0.5$ | $-3.4 \pm 7.2$ | $-0.641$ |
| 48.8 | $18 \pm 0.4$ | $67 \pm 0.04$ | $3.2 \pm 0.5$ | $5.6 \pm 0.2$ | $-8.4 \pm 8.7$ | $-0.659$ |
| 49.6 | $21.9 \pm 0.3$ | $55.6 \pm 0.03$ | $5.5 \pm 0.5$ | $7.1 \pm 0.3$ | $-36.1 \pm 10.9$ | $-0.68$ |
| 50.4 | $23.3 \pm 0.3$ | $52.7 \pm 0.03$ | $8.5 \pm 0.2$ | $9.1 \pm 0.5$ | $-38.2 \pm 11.8$ | $-0.811$ |
| 51.2 | $25.2 \pm 0.3$ | $49.2 \pm 0.03$ | $11.3 \pm 0.5$ | $10.8 \pm 0.4$ | $-45.7 \pm 13.8$ | $-0.885$ |
| 52 | $27 \pm 0.5$ | $46.4 \pm 0.03$ | $15.5 \pm 0.1$ | $11.7 \pm 0.4$ | $-31.1 \pm 21$ | $-0.892$ |

Properties of LD lenses obtained by increasing the width *L* of the simulation box of an almost spherical LD of size *D* = 29.5 *nm* with smallest box width *L* = 47.2 *nm*.

**Table S5:** Properties of symmetrical LD lenses with size *D* = 27.7 *nm*

| $L$ [nm] | $R$ [nm] | $\theta$ [°] | $\Sigma_b$ [mN/m] | $\Sigma_m$ [mN/m] | $\lambda$ [pN] | $\gamma$ [10<sup>9</sup> mN/m<sup>2</sup>] |
| --- | --- | --- | --- | --- | --- | --- |
| 48 | $14.1 \pm 0.4$ | $79.7 \pm 0.1$ | $0.27 \pm 0.15$ | $4.2 \pm 0.5$ | $-2.8 \pm 3.3$ | $-0.901$ |
| 48.8 | $15.9 \pm 0.3$ | $70.1 \pm 0.06$ | $1.5 \pm 0.2$ | $5.2 \pm 0.3$ | $-17.2 \pm 4.3$ | $-0.702$ |
| 49.6 | $18.8 \pm 0.6$ | $59.9 \pm 0.04$ | $3.8 \pm 0.4$ | $6.3 \pm 0.5$ | $-30.3 \pm 10.6$ | $-0.713$ |
| 50.4 | $21.8 \pm 0.5$ | $52.4 \pm 0.04$ | $6.8 \pm 0.3$ | $7.7 \pm 0.4$ | $-35.9 \pm 10$ | $-0.744$ |
| 51.2 | $26.5 \pm 0.7$ | $44.4 \pm 0.03$ | $10.1 \pm 0.3$ | $8.7 \pm 0.3$ | $-36.6 \pm 9.8$ | $-0.687$ |
| 52 | $27.5 \pm 0.4$ | $43 \pm 0.03$ | $13.8 \pm 0.2$ | $11.6 \pm 0.4$ | $-52.9 \pm 11.6$ | $-0.873$ |

Properties of LD lenses obtained by increasing the width *L* of the simulation box of an almost spherical LD of size *D* = 27.7 *nm* with the smallest box width *L* = 48 *nm*.

**Table S6:**
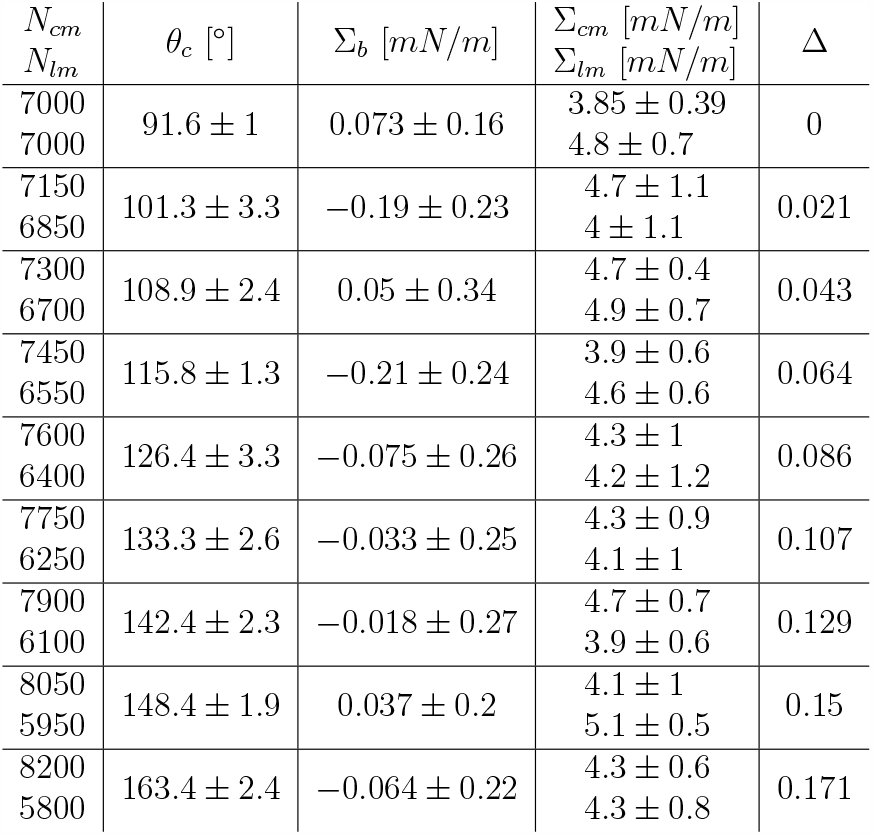
Properties of asymmetric LDs budding to the cytosol

| $N_{cm}$<br>$N_{lm}$ | $\theta_c$ [°] | $\Sigma_b$ [mN/m] | $\Sigma_{cm}$ [mN/m]<br>$\Sigma_{lm}$ [mN/m] | $\Delta$ |
| --- | --- | --- | --- | --- |
| 7000<br>7000 | $91.6 \pm 1$ | $0.073 \pm 0.16$ | $3.85 \pm 0.39$<br>$4.8 \pm 0.7$ | 0 |
| 7150<br>6850 | $101.3 \pm 3.3$ | $-0.19 \pm 0.23$ | $4.7 \pm 1.1$<br>$4 \pm 1.1$ | 0.021 |
| 7300<br>6700 | $108.9 \pm 2.4$ | $0.05 \pm 0.34$ | $4.7 \pm 0.4$<br>$4.9 \pm 0.7$ | 0.043 |
| 7450<br>6550 | $115.8 \pm 1.3$ | $-0.21 \pm 0.24$ | $3.9 \pm 0.6$<br>$4.6 \pm 0.6$ | 0.064 |
| 7600<br>6400 | $126.4 \pm 3.3$ | $-0.075 \pm 0.26$ | $4.3 \pm 1$<br>$4.2 \pm 1.2$ | 0.086 |
| 7750<br>6250 | $133.3 \pm 2.6$ | $-0.033 \pm 0.25$ | $4.3 \pm 0.9$<br>$4.1 \pm 1$ | 0.107 |
| 7900<br>6100 | $142.4 \pm 2.3$ | $-0.018 \pm 0.27$ | $4.7 \pm 0.7$<br>$3.9 \pm 0.6$ | 0.129 |
| 8050<br>5950 | $148.4 \pm 1.9$ | $0.037 \pm 0.2$ | $4.1 \pm 1$<br>$5.1 \pm 0.5$ | 0.15 |
| 8200<br>5800 | $163.4 \pm 2.4$ | $-0.064 \pm 0.22$ | $4.3 \pm 0.6$<br>$4.3 \pm 0.8$ | 0.171 |

Properties of spherical LDs obtained by exchanging lipids from the luminal to the cytosolic monolayer of an initial spherical LD of size *D* = 29.5 *nm* in a fixed simulation box of width *L* = 72 *nm*. The initial spherical LD emerges into a completely budded LD attached to the ER upon exchanging lipids at a fixed total number of lipids *N*_*lip*_ = 14000. *N*_*cm*_ and *N*_*lm*_ denote number of lipids in the cytosolic and luminal leaflets of the LD-monolayers and the ER bilayer. LD angles are calculated based on the LD budding model (Fig. 5b) with *θ*_*l*_ = 180 − *θ*_*c*_.

**Table S7:**
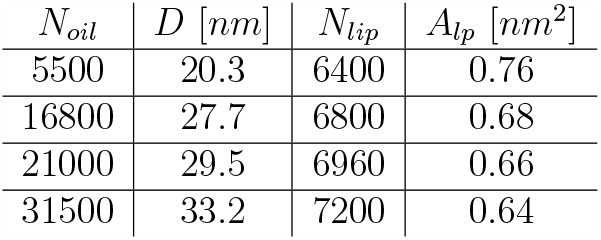
Properties of almost spherical LDs with different sizes

Projected areas-per-lipid *A*_*lp*_ for three nearly-spherical LDs of different sizes.

